# CD4^+^ T cell-derived IL21 regulates stem cell fate in acute myeloid leukemia by activation of p38-MAPK signaling

**DOI:** 10.1101/2023.09.01.555931

**Authors:** Viviana Rubino, Michelle Hüppi, Sabine Höpner, Luigi Tortola, Lea Taylor, Irene Keller, Remy Bruggman, Marie-Noëlle Kronig, Ulrike Bacher, Manfred Kopf, Adrian F. Ochsenbein, Carsten Riether

## Abstract

Self-renewal programs in leukemia stem cells (LSCs) predict poor prognosis in acute myeloid leukemia (AML) patients. We identified CD4^+^ T cell-derived interleukin (IL) 21 as an important negative regulator of self-renewal of murine and human LSCs, but not hematopoietic stem cells. IL21/IL21R signaling favored asymmetric cell division and differentiation in LSCs through accumulation of reactive oxygen species (ROS) and activation of p38-MAPK signaling, resulting in reduced LSCs number and significantly prolonged survival in murine AML models. In human AML, serum IL21 at diagnosis was identified as an independent positive prognostic biomarker for outcome and correlated with better survival and higher complete remission rate in patients that underwent high-dose chemotherapy. IL21 inhibited primary AML LSCs function in vitro by activating ROS and p38-MAPK signaling and this effect was enhanced by cytarabine treatment. Consequently, promoting IL21/IL21R signaling on LSCs may be a novel approach to decrease stemness and increase differentiation in AML.

## Introduction

Acute myeloid leukemia (AML) is an aggressive malignancy that arises from accumulating mutations in hematopoietic precursors of the myeloid lineage. Median age at manifestation of AML is 67 years^1,2^. AML is the most common form of acute leukemia in adults and shows a high mortality^3^. In fact, five-years relative survival rate for AML patients is only 17%, and becomes less than 5% in elderly patients, who have very limited therapeutic options^4,5^. The standard of care for young and fit AML patients consists in intensive chemotherapy, followed by consolidation with chemotherapy or allogeneic hematopoietic stem cells (HSCs) transplantation^5^. In recent years, targeted therapies were introduced and have improved prognosis for distinct genetic subgroups, e.g. fms-related tyrosine kinase 3 gene (*FLT3*)-mutated AML^6^. For patients who cannot tolerate intensive chemotherapy due to comorbidities and/or advanced age, hypomethylating agents combined with the BCL-2 inhibitor Venetoclax became standard of (palliative) AML therapy^7^. Although most patients initially achieve a complete remission (CR) with such treatments, the majority of them will ultimately relapse often with a refractory disease^5^.

Leukemia stem cells (LSCs) are the initiator and driver of the disease. As they escape therapeutic elimination, they are the driver of relapse too. LSCs reside on top of AML hierarchy and are a small pool of primitive cells that unlimitedly self-renew as well as generate the bulk of circulating leukemic blasts^8,9^. Symmetric renewal, defective differentiation and therapy resistance are hallmarks of LSCs that need to be targeted to cure AML^8,10,11^.

LSCs rely for their regulation and maintenance also on interactions with the microenvironment. Similar to their normal HSCs counterpart, LSCs reside in the BM niche, that contains, among others, endothelial cells (ECs), osteoblasts, osteoclasts, adipocytes, mesenchymal stromal cells (MSCs), and nervous fibers^12^. Within the niche, cell-cell contact or cytokines secretion-mediated mechanisms, that originally developed to protect HSCs, are exploited by LSCs to support their own survival and propagation^13,14^. Immune cells are also part of the niche and have been shown to promote leukemia development rather than contributing to leukemia control^15^. However, the interplay between AML cells and adaptive immunity is still poorly understood.

IL21/IL21R signaling is variously involved in immune responses. The IL21R is expressed on several lymphoid and myeloid cell populations and, upon ligation, mainly signals via Janus tyrosine kinases (JAK) and signal transducers and activators of transcription (STAT) and, to a lesser extent, also via phosphoinositol 3-kinase (PI3K)/Akt and mitogen-activated protein kinases (MAPK) pathways^16^. IL21 is primarily produced by activated CD4^+^ T cells and has pleiotropic effects^17–19^. For example, IL21 promotes and sustains cytotoxic CD8^+^ T cell responses during viral infection^20,21^, as well as type-2 CD4^+^ T cell responses during parasitic infection and allergic reactions^19,22,23^. Furthermore, IL21 supports B cell maturation and antibody production^24,25^, suppresses T regulatory cells^19,26^ and enhances expansion and function of natural killer (NK) cells^27,28^. Anti-tumor activity has been widely reported for IL21, in virtue of its role as a mainly immune-activating cytokine^29–33^. Instead, investigations on IL21 direct effect on tumor cells did not yield univocal findings, because of distinct tumor entities and complex tumor-microenvironment interactions. A growth-promoting effect of IL21 has been observed in chronic lymphocytic leukemia^34,35^, follicular lymphoma^36^, Hodgkin’s lymphoma^37^ and multiple myeloma^38^. Conversely, IL21 was shown to have antiproliferative and proapoptotic effects on diffuse large B cell lymphoma^39^. However, if and how IL21/IL21R signaling pathway affects LSCs in AML and whether this knowledge might be translated into clinical application is still unknown.

In this work, we have identified IL21/IL21R signaling pathway as an important regulator of cell fate in human and murine LSCs, but not HSCs. We found that IL21 is a positive prognostic marker for overall survival (OS) and that higher serum IL21 levels correlate with better survival and higher rate of complete remission in patients that undergo high-dose chemotherapy. Our findings therefore suggest that promoting IL21/IL21R signaling on LSCs may be a novel approach to decrease stemness and increase differentiation in AML.

## Results

### Il21/Il21R signaling reduces murine L-GMPs in vivo

We analyzed the expression of the Il21R on leukemic granulocyte-monocyte progenitors (L-GMPs), which represent the LSC population in MLL-AF9 (KMT2A-MLLT3) AML mice, and the level of Il21 in the BM in a murine retroviral transduction/transplantation AML model^40^. L-GMPs expressed the Il21R and IL21 levels were increased in the BM of AML compared to naïve mice (**Figure 1A, B**). To study the effect of Il21 on L-GMPs, we first treated L-GMPs with recombinant mouse (rm)-Il21 followed by plating in methylcellulose. Il21 treatment reduced colony-forming capacity of L-GMPs, suggesting that L-GMPs can directly respond to Il21 (**Figure 1C**).

**Figure 1.**
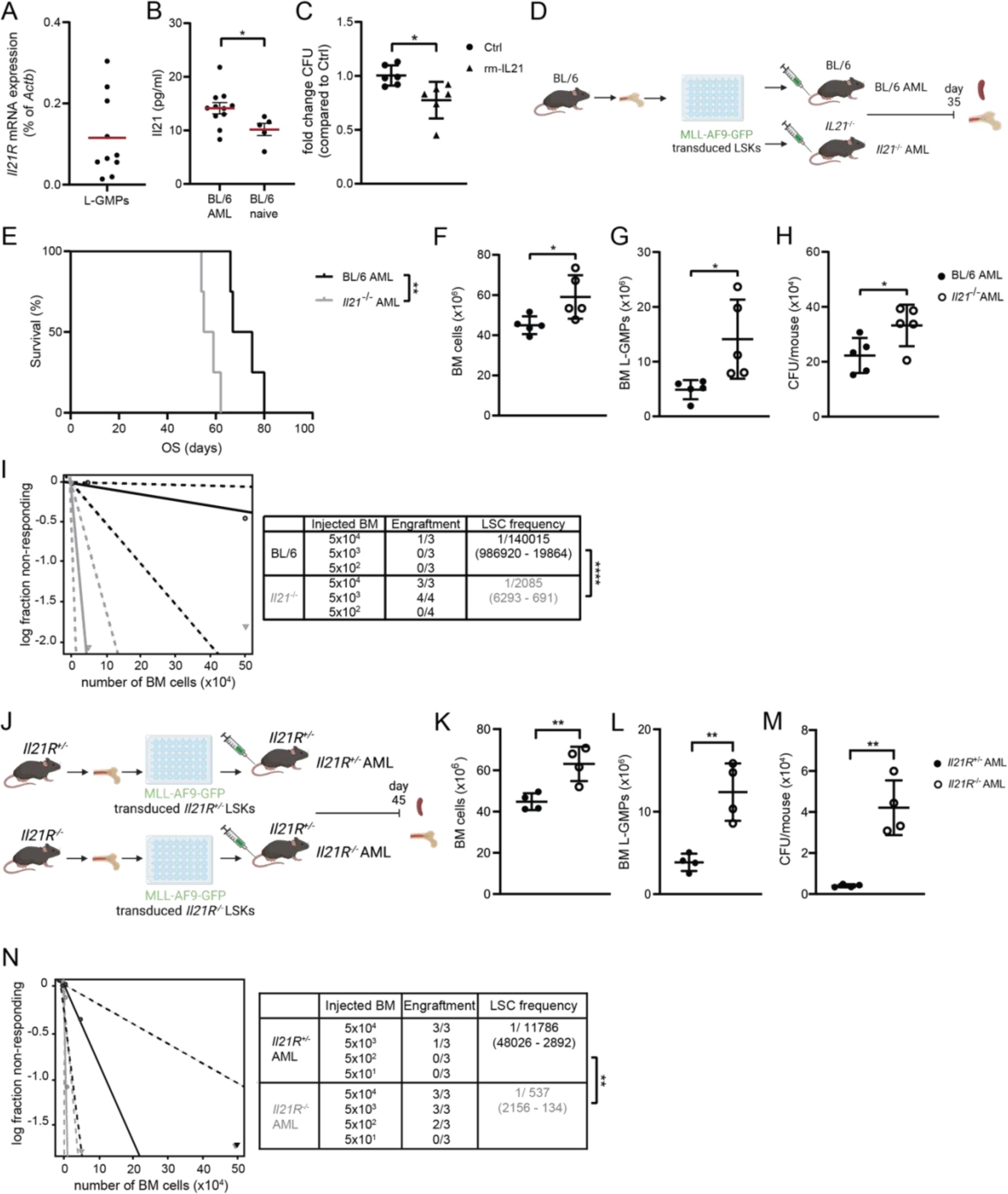
Il21/Il21R signaling reduces murine L-GMPs in vivo. **(A)** *Il21R* mRNA expression (qRT-PCR) in FACS-sorted L-GMPs from the BM of BL/6 AML mice thirty-five days after leukemia transplantation (n = 10). Red bar indicates the mean. **(B)** IL21 levels in BM samples from AML mice (n = 11) and naïve controls (n = 5). Data are shown as mean ± SEM. Statistics were determined by Student’s *t* test. **(C)** Fold change colony-forming units from FACS-sorted L-GMPs cultured in methylcellulose for seven days in the presence or absence of 300 pg/ml rm-IL21 (n= 6 mice/group). Pooled data from two independent experiments are shown and displayed as mean ± SD. Statistics were determined by Student’s *t* test. **(D)** Experimental setup: 5×10^4^ MLL-AF9-GFP-transduced LSKs from the BM of BL/6 donors were injected intravenously into non-irradiated BL/6 and *Il21^-/-^* recipients (BL/6 AML and *Il21^-/-^* AML, respectively). Mice were sacrificed thirty-five days after leukemia transplantation and BM and spleen were analyzed. **(E)** MLL-AF9-GFP AML was induced in BL/6 and *Il21^-/-^*recipients (n = 4 mice/group) and survival was monitored. Statistics were determined by log-rank test. **(F)** BM cellularity and **(G)** number of L-GMPs in BM of BL/6 and *Il21^-/-^* AML mice (n = 5 mice/group). Data are displayed as mean ± SD. Statistics were determined by Student’s *t* test. **(H)** Colony forming units per mouse. 5×10^4^ BM cells were plated into methylcellulose and GFP^+^ colonies were enumerated seven days later by inverted fluorescence microscopy (n = 5 mice/group). Data are displayed as mean ± SD. Statistics were determined by Student’s *t* test. **(F – H)** One representative of four independent experiments is shown. **(I)** Extreme Limiting Dilution Analysis. BM cells from BL/6 and *Il21^-/-^* AML mice were injected at limiting dilutions into lethally irradiated (2 x 6.5 Gy) BL/6 recipients and engraftment was assessed thirty days later. Statistics were determined by χ2 test. **(J)** Experimental setup: 5×10^4^ MLL-AF9-GFP-transduced LSKs from the BM of *Il21R^+/-^* and *Il21R^-/-^* donors were injected intravenously into non-irradiated *Il21R^+/-^* recipients (*Il21R^+/-^* and *Il21R^-/-^*AML, respectively). Mice were sacrificed forty-five days after leukemia transplantation and BM and spleen were analyzed. **(K)** BM cellularity, **(L)** number of L-GMPs in BM and **(M)** colony forming units per mouse (n = 4 mice/group). Data are displayed as mean ± SD. Statistics were determined by Student’s *t* test. **(N)** Extreme Limiting Dilution Analysis. BM cells from *Il21R^+/-^* and *Il21R^-/-^* AML mice were injected at limiting dilutions into lethally irradiated (2 x 6.5 Gy) BL/6 recipients and engraftment was assessed thirty days later. Statistics were determined by χ2 test. **(K– M)** One representative of two independent experiments is shown. *, P < 0.05; **, P < 0.01; ****, P < 0.0001. Abbreviations: L-GMPs, leukemic granulocyte-macrophage progenitors; LSKs, Lin^-^ Sca-1^+^ c-kit^+^; Lin, lineage; OS, overall survival; BM, bone marrow; CFU, colony-forming units; LSC, leukemic stem cell. See also Figure S1 and S2.

To study the functional relevance of Il21/Il21R signaling on AML development in vivo, we transplanted MLL-AF9-GFP-transduced BL/6 lineage^-^Sca-1^+^c-kit^+^ cells (LSKs) into non-irradiated BL/6 mice and *Il21*^-/-^ mice (BL/6 AML and *Il21*^-/-^ AML, respectively) (**Figure 1D**). AML development was faster and resulted in significantly shorter survival for *Il21*^-/-^ AML mice compared to BL/6 AML mice **(Figure S1A and Figure 1E**). To determine the role of Il21/Il21R signaling on L-GMPs, BL/6 and *Il21*^-/-^ AML mice were sacrificed thirty-five days after AML induction and spleen and BM were analyzed. Leukemia burden as indicated by BM cellularity, numbers of MLL-AF9-GFP^+^Gr1^+^Cd11b^+^ leukemic cells in the spleen and in the BM and leukemic blasts frequency in the BM was smaller in BL/6 AML mice compared to *Il21*^-/-^ AML mice (**Figure 1F and Figures S1B-E)**. In addition, the frequency of primitive AML cells (MLL-AF9-GFP^+^lin^-^ cells) was substantially reduced in BL/6 AML mice **(Figure S1F).** Similarly, the number of L-GMPs was significantly diminished in BL/6 AML mice phenotypically and functionally, as assessed by FACS and colony formation assays (**Figures 1G, H**).

The most stringent way to address changes in stem cell frequencies is to transplant whole BM cells at titrated numbers into lethally irradiated mice^41^. To functionally investigate leukemia-initiating cells in vivo, we transferred BM cells from primary BL/6 and *Il21*^-/-^ AML mice at titrated numbers into lethally irradiated secondary recipients. Extreme limiting-dilution analysis (ELDA)^41^ revealed that the presence of Il21 substantially reduced the frequency of L-GMPs in limiting-dilution experiments in vivo by a factor of 67 (**Figure 1I**).

Comparable results on L-GMPs and AML development have been obtained when MLL-AF9-GFP-transduced and MLL-ENL-YFP-transduced *Il21R*^+/-^ and *Il21R*^-/-^ LSKs were transplanted into *Il21R*^+/-^ control mice (*Il21R*^+/-^ AML and *Il21R*^-/-^ AML, respectively) (**Figures 1J-N and Figures S1G-I and S2A-E**), indicating that IL21/IL21R signaling on AML cells regulates leukemogenesis.

Importantly, *Il21R* deficiency on LSKs did not affect their repopulating capacity in steady-state and stress-induced hematopoiesis. **(Figure S2F, G)**.

In summary, these data suggest that *Il21/Il21R* signaling affects stem cell function in AML but not in normal and demand-adapted hematopoiesis.

### Il21/Il21R signaling reduces stem cell maintenance and triggers differentiation-promoting signaling pathways in L-GMPs

To analyze the molecular mechanism of how IL21 affects stemness of L-GMPs, we performed bulk RNA-seq analysis on L-GMPs derived from the BM of BL/6 AML and *Il21*^-/-^ AML mice. Overall, 72 genes were differentially expressed between BL/6 and *Il21*^-/-^ L-GMPs, with 44 and 28 genes being up- and down-regulated, respectively (**Figure 2A**). Gene ontology (GO) and gene set enrichment analysis (GSEA) revealed that Il21/Il21R signaling in L-GMPs reduced gene expression signatures related to proliferation, reactive oxygen species (ROS) production, mitochondrial activity and stemness, as well as stem cell-related signaling pathways such as WNT, NF-kB and MAPK signaling, and promoted differentiation signatures **(Figure 2B, C)**.

**Figure 2.**
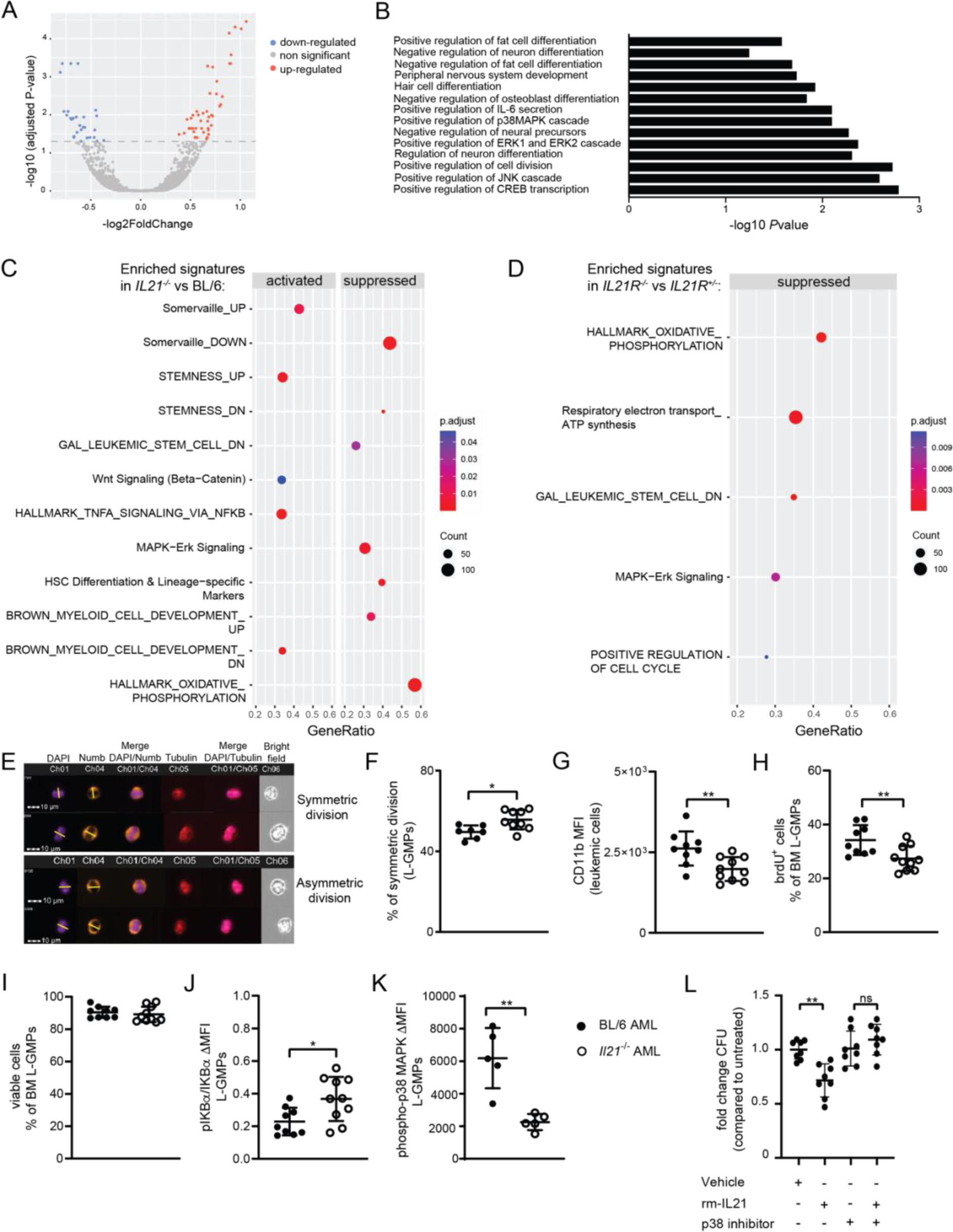
Il21/Il21R signaling reduces stem cell-related signaling pathways and triggers differentiation-promoting signaling pathways in L-GMPs. **(A)** Volcano plot of differentially expressed genes in L-GMPs from BM of BL/6 and *Il21^-/-^* AML mice (n= 3 mice/group). Log2 fold differences of gene expression levels in L-GMPs from BM of *Il21^-/-^* AML mice versus L-GMPs from BM of BL/6 AML mice are shown. **(B)** Bar plot for the -log10 of the p-value of selected GO terms (biological process), showing enriched pathways of differentially expressed genes. **(C, D)** Gene set enrichment analysis (GSEA) showed the activated and suppressed pathways in **(C)** L-GMPs from *Il21^-/-^*AML mice versus L-GMPs from BL/6 AML mice and in **(D)** L-GMPs from *Il21R^-/-^*AML mice versus L-GMPs from *Il21R^+/-^* AML mice (n=3 mice/group). A dot plot was generated to show the most significant enriched terms, with dot size indicating gene counts and dot color indicating the enrichment scores as adjusted p-values. **(E)** Representative picture of Numb distribution in dividing FACS-purified L-GMPs from BM of BL/6 and *Il21^-/-^*AML mice. Cells were analyzed by ImageStream^®^xMkII. Nuclei are stained with DAPI (in violet), α-tubulin is stained in red and Numb in orange. Cell division plane (yellow line) was assigned based on α-tubulin and the cleavage furrow. **(F)** Quantification of L-GMPs from BM of n = 7 BL/6 and n = 9 *Il21^-/-^* AML mice in symmetric cell division. Statistics were determined by Student’s *t* test. **(E, F)** Two pooled independent experiments are shown (n = 3 - 5 mice/group). **(G)** CD11b mean fluorescence intensity of MLL-AF9-GFP^+^ leukemic cells from BM of BL/6 and *Il21^-/-^* AML mice (n= 9-10 mice/group). **(H)** Quantification of proliferating L-GMPs measured as brdU incorporation in vivo, **(I)** cell viability measured as percentage of AnnexinV^-^ L-GMPs, (**J)** NF-kB pathway activation measured as ratio between protein expression of IκBα and its phosphorylated form pIκBα in L-GMPs from BM of BL/6 and *Il21^-/-^* AML mice (n= 9-10 mice/group). **(G - J)** Two pooled independent experiments are shown (n = 5 - 6 mice/group). **(K)** delta of the geometric mean fluorescence intensity (MFI) of phospo-p38 MAPK staining versus its isotype control on L-GMPs from BM of BL/6 and *Il21^-/-^* AML mice (n = 5 mice/group). **(L)** FACS-purified L-GMPs from BL/6 mice were pre-treated with 10 nm/ml of the p38 MAPK inhibitor SB203580 or vehicle prior to overnight culture in in the presence or absence of 300 pg/ml rm-IL21 and plated in methylcellulose. Fold-change in colony formation is depicted in the graph. Pooled data from two independent experiments are shown and displayed as mean ± SD. Statistics were determined by Student’s *t* test. *, P < 0.05; **, P < 0.01. Abbreviations: PC, principal component; L-GMPs, leukemic granulocyte-macrophage progenitors; brdU, bromodeoxyuridine. See also Figure S3.

To identify the shared molecular mechanisms between L-GMPs derived from a IL21-deficient microenvironment and L-GMPs lacking the IL21R, we next performed bulk RNA-seq analysis on L-GMPs derived from *Il21R*^+/-^ AML and *Il21R*^-/-^ AML mice. Similar to L-GMPs from *Il21*^-/-^ AML mice, *Il21R^-/-^* L-GMPs showed alterations in gene expression signatures related to stemness, ROS production, mitochondrial activity and proliferation (**Figure 2D**).

### Il21/Il21R signaling reduces stemness of L-GMPs by favoring asymmetric division and activating the p38-MAPK signaling pathway

To confirm that Il21/Il21R signaling reduces stemness and favors differentiation potential in AML, we first addressed asymmetric division and symmetric renewal of L-GMPs and CD11b expression of bulk leukemia cells. We found that L-GMPs in the BM of *Il21*^-/-^ AML mice showed higher symmetric division rate compared to BL/6 AML controls (**Figure 2E, F**), suggesting that *Il21/Il21R* signaling regulates differentiation of L-GMPs through promotion of asymmetric cell division over symmetric renewal. In line with these findings, bulk leukemia cells in the BM of *Il21*^-/-^ AML mice had a lower expression of the differentiation marker CD11b compared to bulk BL/6 leukemia cells (**Figure 2G**). In addition, L-GMPs in the BM of *Il21*^-/-^ AML mice diluted BrdU significantly more compared to controls in 48h label-retaining experiments in vivo which is indicative of a rapidly dividing L-GMP population. (**Figure 2H**). Furthermore, L-GMPs in the BM of *Il21*^-/-^ AML mice, in spite of identical cell viability, showed increased phosphorylation of IKBα and decreased phosphorylation of p38-MAPK, indicative of altered NF-kB and p38-MAPK signaling activity (**Figure 2I-K and Figure S3A)**. AML stem cell properties can also be defined based on ROS levels and mitochondrial dynamics. Consistent with a less differentiated AML phenotype, staining of intracellular ROS with the cell-permeant dye CellRox^TM^ revealed a smaller percentage of CellRox^+^ L-GMPs in the BM of *Il21*^-/-^ AML mice compared to BL/6 AML mice **(Figure S3B)**. In addition, staining with the mitochondrial probes MitoTracker^TM^ and TMRM^TM^ revealed respectively that L-GMPs in the BM of *Il21*^-/-^ AML mice had increased mitochondrial mass, without displaying changes in their mitochondrial membrane potential, which is consistent with LSCs dependency on oxidative phosphorylation (OXPHOS) for their metabolism **(Figure S3C, D)**.

To verify that the cellular processes and signaling cascades identified in *Il21^-/-^* AML mice are also modulated in *Il21R^-/-^*AML mice, we assessed CD11b expression on bulk leukemia cells as well as phosphorylation states of IKBα and p38-MAPK in L-GMPs from *Il21R*^-/-^ and control *Il21R*^+/-^AML mice. In line with the findings obtained in *Il21^-/-^* AML mice, *Il21R*^-/-^ AML mice had a lower expression of CD11b on bulk leukemia cells **(Figures S3E)**. Furthermore, phosphorylation of p38-MAPK in *Il21R^-/-^* L-GMPs was significantly reduced compared to controls, while phosphorylation of IKBα was unchanged between the groups **(Figures S3F-H)**.

To functionally demonstrate that the IL21-mediated regulation of L-GMPs is dependent on p38-MAPK signaling, we incubated L-GMPs with recombinant mouse (rm)-Il21 after pre-incubation with the potent p38-MAPK inhibitor SB203580 that does not compromise the activity of the ERK or JNK MAP kinase subgroups^42^ and assessed colony formation in vitro. Effect of Il21 on colony formation could be reverted by blockade of p38-MAPK signaling (**Figure 2L**).

Overall, these findings suggest that *Il21/Il21R* signaling regulates cell fate of L-GMPs in AML by inducing differentiation, accumulation of ROS and activation of the p38-MAPK signaling pathway.

### CD4^+^ T cell-derived Il21 reduces stemness of murine LSCs in vivo

To determine the source of Il21 in MLL-AF9 AML mice, we induced MLL-AF9 AML in *Il21*^mcherry^ reporter mice^43^. CD4^+^ T cells were identified as the primary source of IL21 in BM, blood and spleen of MLL-AF9 AML mice by FACS (**Figure 3A, B and data not shown)**. However, no difference in the frequency of Il21-producing CD4^+^ T cells was observed between naïve and AML mice. Similar results on Il21 production by CD4^+^ T cells were obtained by qRT-PCR **(Figure S4A)**. L-GMPs and CD45^-^lineage^-^ (CD45^-^lin^-^) BM cells which comprise classical niche cells in AML such as MSCs and ECs did not express IL21 mRNA **(Figure S4B and data not shown)**.

**Figure 3.**
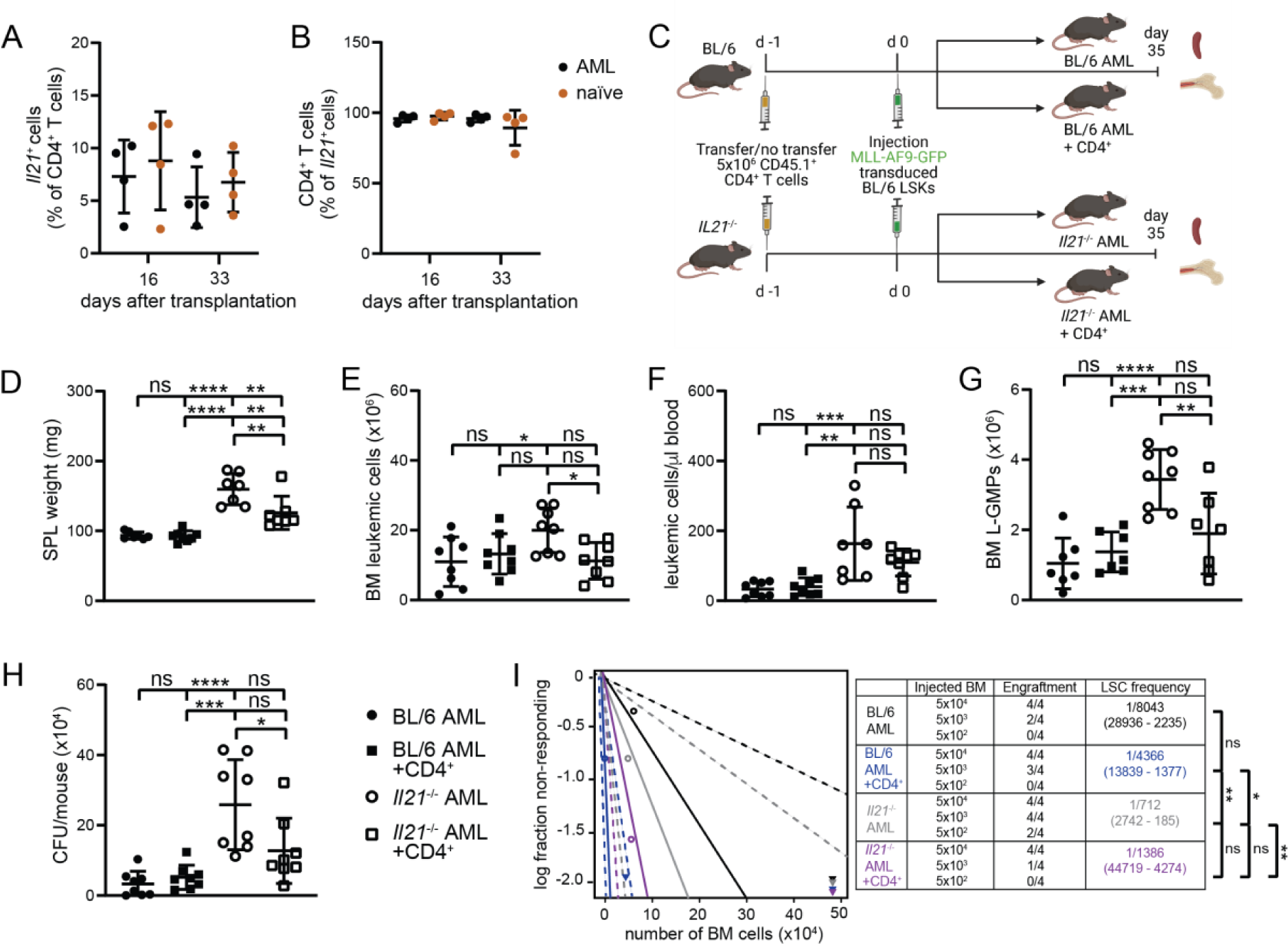
CD4^+^ T cell-derived Il21 reduces murine LSCs in vivo. **(A, B)** MLL-AF9 AML was induced in *Il21^mcherry^* reporter mice. Frequencies of *mcherry-Il21*^+^ CD4^+^ T cells **(A)** and CD4^+^ *mcherry-Il21*^+^ cells **(B)** were determined by flow cytometry 16 and 33 days after leukemia transplantation, in BM of AML and naïve *Il21^mcherry^* mice (n = 4 mice/group). **(C)** Experimental setup: 5×10^6^ CD4^+^ T cells were FACS-sorted from the spleens of CD45.1 mice and injected intravenously into two out of four experimental groups (non-irradiated BL/6 and *Il21^-/-^* recipients) one day prior to leukemia transplantation. One day after, 5×10^4^ MLL-AF9-GFP-transduced LSKs from the BM of BL/6 donors were injected intravenously into all four experimental groups (BL/6 AML, *Il21*^-/-^ AML, BL/6 AML + CD4^+^ and *Il21*^-/-^ AML + CD4^+^, respectively). Mice were sacrificed 35 days after leukemia transplantation and BM and spleen were analyzed (n = 4 mice/group). **(D)** Spleen size, **(E)** number of MLL-AF9-GFP^+^ leukemic cells in BM and **(F)** peripheral blood, **(G)** number of L-GMPs in the BM of BL/6 AML, *Il21*^-/-^ AML, BL/6 AML + CD4^+^ and *Il21*^-/-^ AML + CD4^+^ mice 35 days after leukemia transplantation. **(H)** Colony forming units per mouse. 5×10^4^ BM cells were plated into methylcellulose and GFP^+^ colonies were enumerated seven days later by inverted fluorescence microscopy. **(D - H)** Two pooled independent experiments are shown (n = 4 mice/group/experiment). Data are shown as mean ± SD. Statistics were determined by one-way ANOVA followed by Tukey’s multiple comparisons test. **(I)** Extreme Limiting Dilution Analysis. BM cells from BL/6 AML, *Il21*^-/-^ AML, BL/6 AML + CD4^+^ and *Il21*^-/-^ AML + CD4^+^ mice were injected at limiting dilutions into lethally irradiated (2 x 6.5 Gy) BL/6 recipients and engraftment was assessed thirty days later. Statistics were determined by χ2 test. *, P < 0.05; **, P < 0.01; ***, P < 0.001; ****, P < 0.0001. Abbreviations: LSKs, Lin^-^ Sca-1^+^ c-kit^+^; SPL, spleen; L-GMPs, leukemic granulocyte-macrophage progenitors; CFU, colony-forming units. See also Figure S4.

To demonstrate that Il21 derived from CD4^+^ T cells contributes to AML development, we adoptively transferred congenic CD45.1^+^ Il21-proficient CD4^+^ T cells into BL/6 AML and *Il21*^-/-^ AML mice (**Figure 3C**). Importantly, at the time of analysis, adoptively transferred CD4^+^ T cells could be detected in BM, spleen and PB of BL/6 AML and *Il21*^-/-^ AML mice **(Figure S4C)**. Transfer of CD4^+^ T cells into *Il21*^-/-^ AML significantly reduced leukemia burden and L-GMP numbers and frequency to levels comparable to BL/6 AML mice (**Figures 3D - I**). In summary, these findings suggest that CD4^+^ T cell-derived Il21 inhibits leukemia development and stemness in AML.

### IL21 levels are elevated in the serum of AML patients at diagnosis and are a positive prognostic marker for OS

To analyze the role of IL21 in human AML, we first determined IL21 levels in the serum (sIL21) of newly diagnosed AML patients **(Table S1)**. Because AML is primarily a disease of the elderly population, we initially verified in a publicly available resource^44^ that sIL21 levels are not altered with age in healthy individuals **(Figure S5A).** In contrast, sIL21 levels were significantly increased in 193 AML patients compared to age-matched healthy controls (**Figure 4A**, mean sIL21 81.6 and 7.9 pg/ml, respectively). Kaplan-Meier analysis revealed that patients with high levels of sIL21 (≥ 29 pg/ml) survived substantially longer than patients with low levels of sIL21 (**Figure 4B**). A similar effect of sIL21 levels on OS was obtained when the patient cohort was subdivided in patients with low, intermediate, and high levels of sIL21 **(Figure S5B).** Well-established risk factors for OS in AML such as patients’ age and cytogenetic/molecular risk group^5^ did not act as confounding factors in our analysis (**Figure 4C, D**). Similarly, sIL21 did not correlate with the numbers of CD4^+^ T cells in blood, which has been identified as a primary source of IL21 **(Figure S5C)**. Furthermore, multivariate analysis for sIL21 levels adjusted for patient age, risk group, blast percentage in blood and BM as well as leukocytes counts substantiated sIL21 as an independent positive prognostic marker in AML (**Figure 4E**). Similar to sIL21, high levels of *IL21* mRNA were associated with a favorable prognosis in two independent AML microarray datasets^45,46^ **(Figure 4F, G)**. Like for sIl21 levels, *IL21* mRNA did not correlate with *CD4* mRNA levels in either dataset **(Figures S5D, E)**. Overall, these results identify sIL21 and *IL21* mRNA as independent positive prognostic biomarkers for OS in AML.

**Figure 4.**
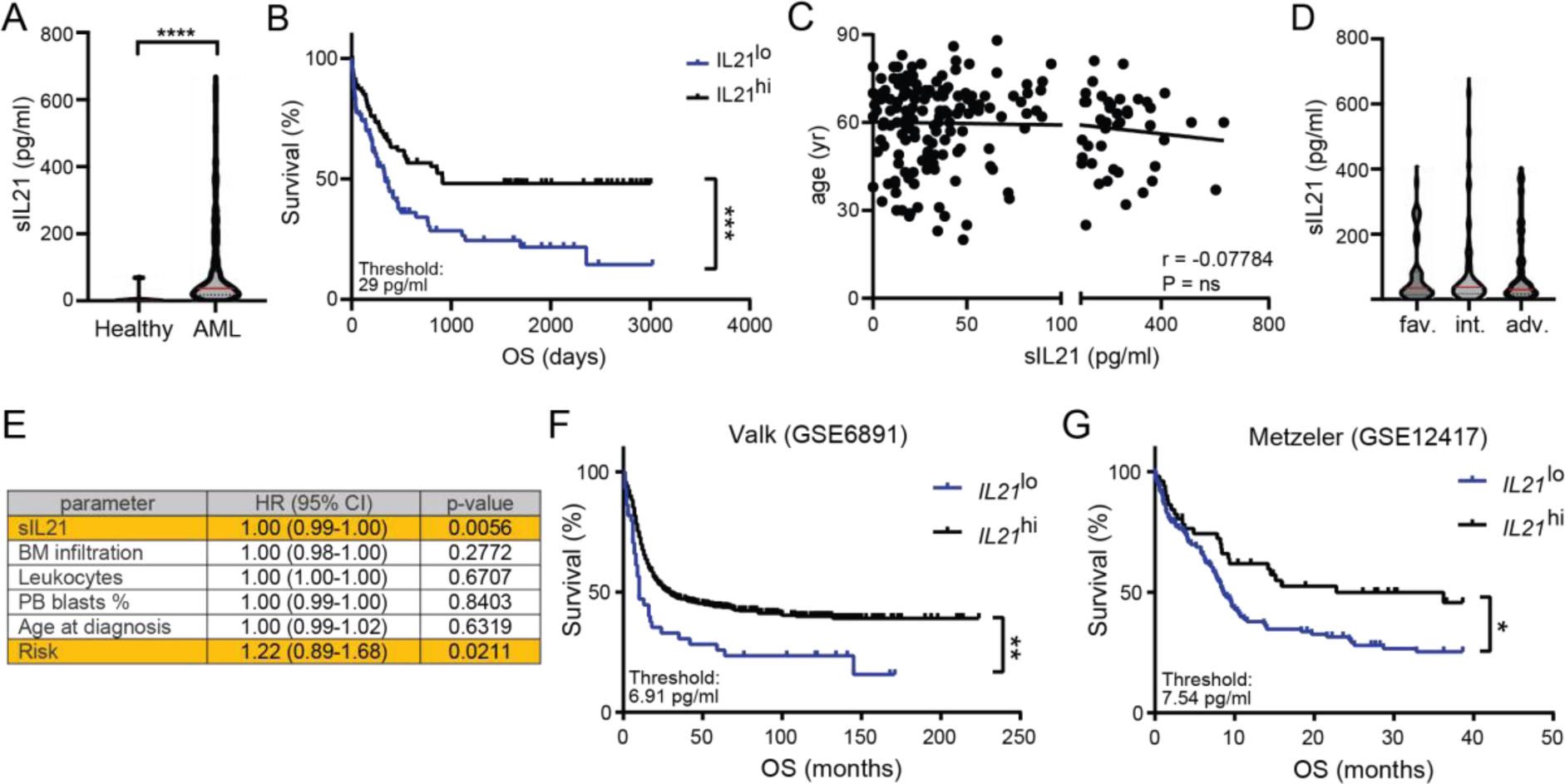
IL21 is an independent positive prognostic marker for OS in AML. **(A)** IL21 levels in serum samples from newly diagnosed AML patients (n = 193) and age-matched healthy controls (n = 10). Red bars indicate the mean. Statistics were determined by Mann-Whitney test. **(B)** Kaplan-Meier survival curves of the AML patients’ cohort (n = 193) divided into two groups at the IL21 serum levels (sIL21) threshold of 29 pg/ml. Statistics were determined by log-rank test. **(C)** Correlation of patients’ age with sIL21 levels. Statistics were determined by Pearson r test. **(D)** sIL21 of patients in the different risk groups. Red bars indicate the mean. Statistics were determined by one-way ANOVA. **(E)** Multivariate analysis for sIL21 adjusted for BM infiltration, leukocyte counts, percentage of peripheral blood blasts, age and risk group. Statistics were determined by multiple Cox regression. **(F, G)** Two publicly available microarray datasets were analyzed for *IL21* mRNA expression levels and their association with prognosis. Statistics were determined by log-rank test. **(F)** Valk dataset, accession number <u>GSE6891</u>, sIL21 threshold 6.91 pg/ml. **(G)** Metzeler dataset, accession number <u>GSE12417</u>, sIL21 threshold 7.54 pg/ml. *, P < 0.05; **, P < 0.01; ***, P < 0.001; ****, P < 0.0001. Abbreviations: OS, overall survival; yr, years; fav., favorable; int., intermediate; adv., adverse; HR, hazard ratio; CI, confidence interval; PB, peripheral blood. See also Figure S5.

### AML stem and progenitor cells but not normal hematopoietic stem and progenitor cells express the IL21 receptor

We analyzed the source of IL21 in AML by qRT-PCR. Similar to the data obtained in the murine AML model, *IL21* mRNA was mostly expressed by CD4^+^ T cells (22 out of 32 patients analyzed) but not CD8^+^ T cells and CD34^+^ AML stem and progenitor cells (LSPCs) in the BM of newly diagnosed AML patients (**Figure 5A and Figure S6A** for LSPCs gating strategy). Next we determined the expression of the IL21R and its co-receptor CD132 on AML LSPCs. In human AML, the IL21R was expressed on T cells but also on LSPCs in 21/35 BM samples and 12/30 blood samples by FACS and qRT-PCR (**Figure 5B-D, Figure S6B)**. CD132, the co-receptor for IL21R, was expressed on LSPCs from all patients analyzed (data not shown). *IL21R* mRNA expression on LSPCs could not be associated with *IL21* mRNA expression by CD4^+^ T cells (**Figure 5E**). Importantly, normal hematopoietic stem and progenitor cells (HSPCs) derived from the BM of healthy donors did not express the IL21R on the cell surface (**Figure 5F and Figure S6C** for HSPCs gating strategy). To analyze the expression of the IL21R during emergency hematopoiesis in humans, we stained the IL21R by FACS in 15 multiple myeloma patients which underwent allogeneic stem cells transplantation. The vast majority of HSPCs (12/15) derived from the BM of these patients 14 days after stem cell transplantation did not express the IL21R on the surface (**Figure 5G**). These results indicate that stem and progenitor cells from the BM of most AML patients express the IL21R and that CD4^+^ T cells are the primary source of IL21 in human AML.

**Figure 5.**
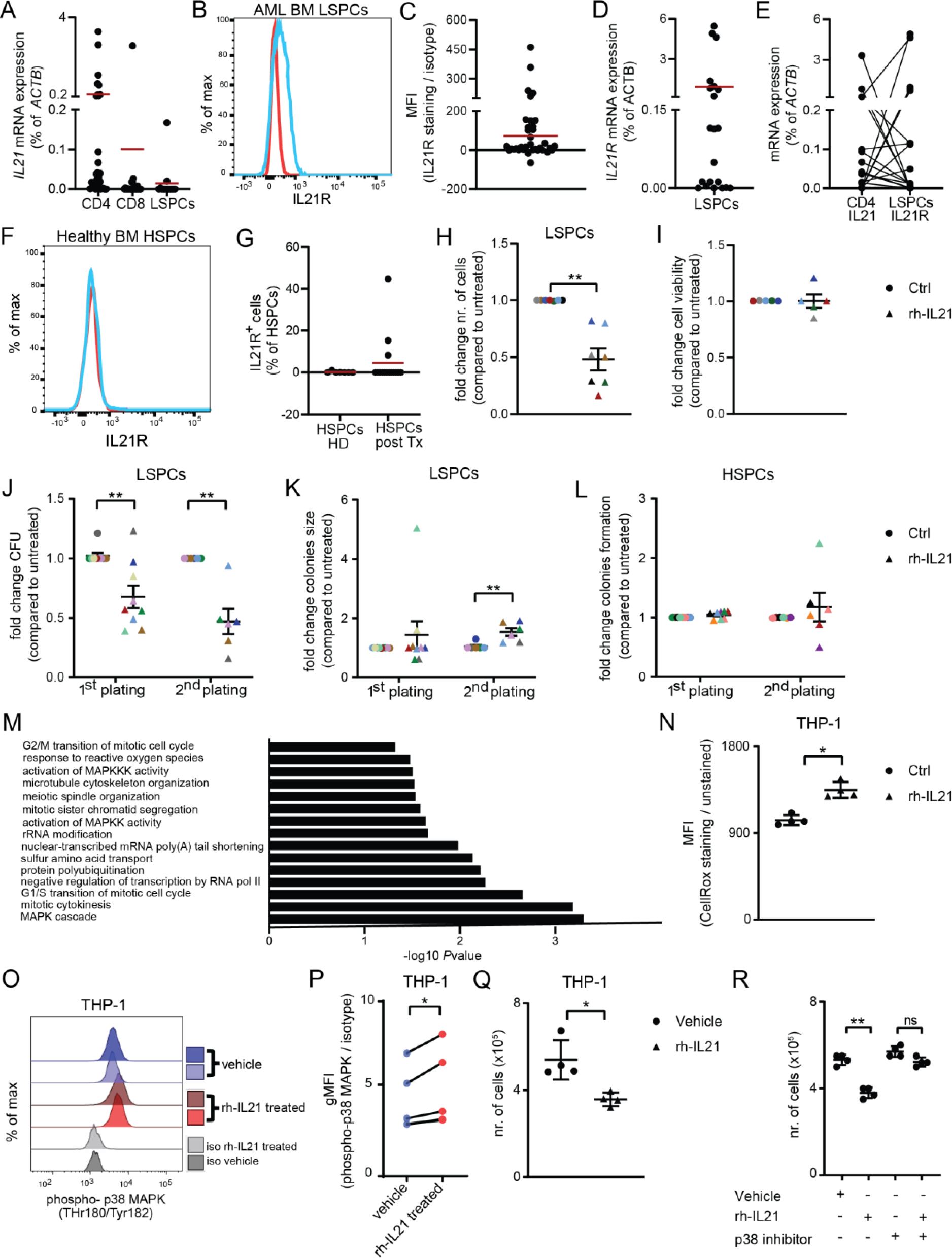
IL21 is produced by CD4^+^ T cells and reduces cell growth and colony-forming capacity of AML stem and progenitor cells in vitro. **(A**) *IL21* mRNA expression (qRT-PCR) in FACS-sorted CD4^+^ T cells (n = 32), CD8^+^ T cells (n = 27) and CD45^dim^SSC^lo^lin^-^CD34^+^ LSPCs (n = 13) from newly diagnosed AML patients. Red bars indicate the mean. Statistics were determined by one-way ANOVA. **(B)** Representative histogram of IL21R (blue line) and relative isotype control (red line) staining on BM LSPCs. **(C)** Mean fluorescence intensity (MFI) quotient of IL21R staining versus its isotype control on LSPCs (n = 35) from BM samples of newly diagnosed AML patients. Red bar indicates the mean. (D) *IL21R* mRNA expression (qRT-PCR) in FACS-sorted LSPCs from newly diagnosed AML patients (n = 21). Red bar indicates the mean. (E) *IL21* and *IL21R* mRNA expression (qRT-PCR) in paired CD4^+^ T cells and LSPCs FACS-sorted from newly diagnosed AML patients (n = 21). **(F)** Representative histogram of IL21R (blue line) and relative isotype control (red line) staining on BM HSPCs of healthy controls. **(G)** Percentage of HSPCs from the BM of healthy donors (n = 7) and multiple myeloma patients who underwent allogeneic hematopoietic stem cells transplantation (n = 15) that express IL21R. Red bars indicate the mean. **(H)** Cell number (n = 7) and **(I)** viability (n = 5) of FACS-sorted LSPCs cultured in vitro for 72 h in the presence or absence of 100 pg/ml rh-IL21. **(J)** LSPCs colonies and **(K)** number of cells per colony were enumerated after two weeks of culture in methylcellulose in the presence or absence of 100 pg/ml rhIL21 (two rounds of plating, n = 9). **(L)** HSPCs colonies were enumerated after two weeks of culture in methylcellulose in the presence or absence of 100 pg/ml rh-IL21 (1^st^ plating, n = 8; 2^nd^ plating, n = 6). **(H - L)** Each dot represents the mean of three technical replicates. Different colors indicate different patients. Statistics were determined by Student’s *t* test. Data are shown as mean ± SEM. **(M)** FACS-sorted CD45^dim^SSC^lo^lin^-^CD34^+^ LSPCs from three AML patients were cultured in vitro in the presence or absence of 100 pg/ml rh-IL21. After 72h of culture, RNA was extracted and sequenced. The bar plot for the -log10 of the p-value of selected GO terms shows enriched pathways of differentially expressed genes. **(N)** Intracellular reactive oxygen species measured by CellRox^TM^ staining, **(O)** histograms showing phosphorylation of p38 MAPK (phospho-p38 MAPK), **(P)** delta of the geometric mean fluorescence intensity (MFI) of phospo-p38 MAPK staining versus its isotype control and **(N)** number of THP-1 cells cultured in vitro for 72 h in the presence or absence of 1 nm/ml rh-IL21. Pooled data from four independent experiments are shown, with each dot representing the mean of three to five technical replicates. Statistics were determined by paired Student’s *t* test. **(R)** Number of THP-1 cells untreated or pretreated with10 nm/ml of the p38 MAPK inhibitor SB203580 were cultured in vitro for 72 h in the presence or absence of 1 ng/ml rh-IL21 Pooled data from four independent experiments are shown and each dot represents the mean of three technical replicates. Data are shown as mean ± SD. Statistics were determined by Student’s *t* test. *, P < 0.05; **, P < 0.01; ****, P < 0.0001. Abbreviations: LSPCs, leukemic stem and progenitor cells; MFI, mean fluorescence intensity; HSPCs, hematopoietic stem and progenitor cells; HD, healthy donor; Tx, transplantation. See also Figure S6.

### IL21/IL21R signaling on LSPCs inhibits cell growth and stemness in vitro through activation of p38-MAPK signaling

To study the functional role of IL21/IL21R signaling in AML stem and progenitor cells, FACS-sorted CD45^dim^SSC^lo^lin^-^CD34^+^ LSPCs from different cytogenetic/molecular risk groups were cultured in the presence of 100 pg/ml of recombinant human (rh)-IL21 for 72 hours. 100 pg/ml of rh-IL21was selected because it resembles the mean concentration of IL21 detected in serum of AML patients (**Figure 4A**). Addition of rh-IL21 to the culture significantly reduced cell numbers per well without affecting cell viability (**Figure 5H, I**).

To assess colony-forming capacity, LSPCs were cultured overnight in presence and absence of rh-IL21, followed by plating in methylcellulose. rh-IL21 treatment significantly reduced colony formation of LSPCs (**Figure 5J**). Replating revealed that this effect was maintained even in the absence of rh-IL21 (**Figure 5J**). In addition, rh-IL21 treatment resulted in increased colony size after replating (**Figure 5K**). In contrast, rh-IL21 treatment did not affect the clonogenic potential of HSPCs from healthy BM donors (**Figure 5L**).

In our murine AML models, the p38-MAPK signaling was identified as the central pathway that regulates stemness of L-GMPs (**Figure 2 and Figure S3)**. To verify that p38-MAPK signaling is also active in primary human LPSCs, we incubated CD34^+^ LSPCs from 3 newly diagnosed AML patients **(Table S2)** in presence and absence of rh-IL21 and performed bulk RNA-seq. GO analysis revealed that IL21 triggered pathways related to cell cycle, ROS and MAPK signaling (**Figure 5M**).

To mechanistically confirm that exposure to rhIL-21 promotes the accumulation of ROS and activation of the p38-MAPK signaling pathway in human AML cells, we incubated the human THP-1 AML cells with rh-IL21 for 72 hours. THP-1 AML cells have been previously shown to express the IL21R on the cell surface^47^. Culture of THP-1 AML cells in presence of rh-IL21 increased cellular ROS and p38-MAPK levels resulting in reduced cell growth (**Figure 5N-Q**). Blockade of p38-MAPK signaling with the p38-MAPK inhibitor SB203580 restored growth of THP-1 AML cells almost to the level of vehicle-treated THP-1 AML cells (**Figure 5R**). These findings indicate that the IL21/IL21R interaction reduces cell growth and self-renewal of LSPCs, but not of normal HSPCs, through accumulation of ROS and activation of p38-MAPK signaling.

### IL21/IL21R signaling promotes the sensitivity of AML LSCs to cytarabine treatment

Our functional and transcriptomic analysis revealed that activation of IL21/IL21R signaling reduces murine and human LSPCs stemness. We therefore hypothesized that IL21/IL21R signaling might render LSPCs more susceptible to chemotherapy. To test this hypothesis, we first analyzed sIL21 levels only in 110 patients that received intensive chemotherapy (“7+3” regimen) as a first line treatment. Importantly, patients that achieved CR after induction chemotherapy survived substantially longer compared to patients that did not achieve CR (**Figure 6A**). Kaplan-Meier analysis of this cohort further revealed that patients with high levels of sIL21 (≥ 72 pg/ml) survived significantly longer than patients with low and intermediate levels of sIL21 (**Figure 6B**). Similarly, we found that sIL21 levels are increased in patients at diagnosis that achieved CR compared to patients that did not achieve CR (**Figure 6C**). Patients’ age and cytogenetic/molecular risk category did not act as confounding factors in our analysis **(Figure S7A, B)**. Furthermore, CR rate was found to be significantly higher in the subgroups with high and intermediate sIL21 levels compared to the subgroup with low sIL21 levels (81% versus 59%) (**Figure 6D**). In contrast, sIL21 levels at diagnosis did not correlate with OS of patients that received first line palliative treatment **(Figure S7C)**.

**Figure 6.**
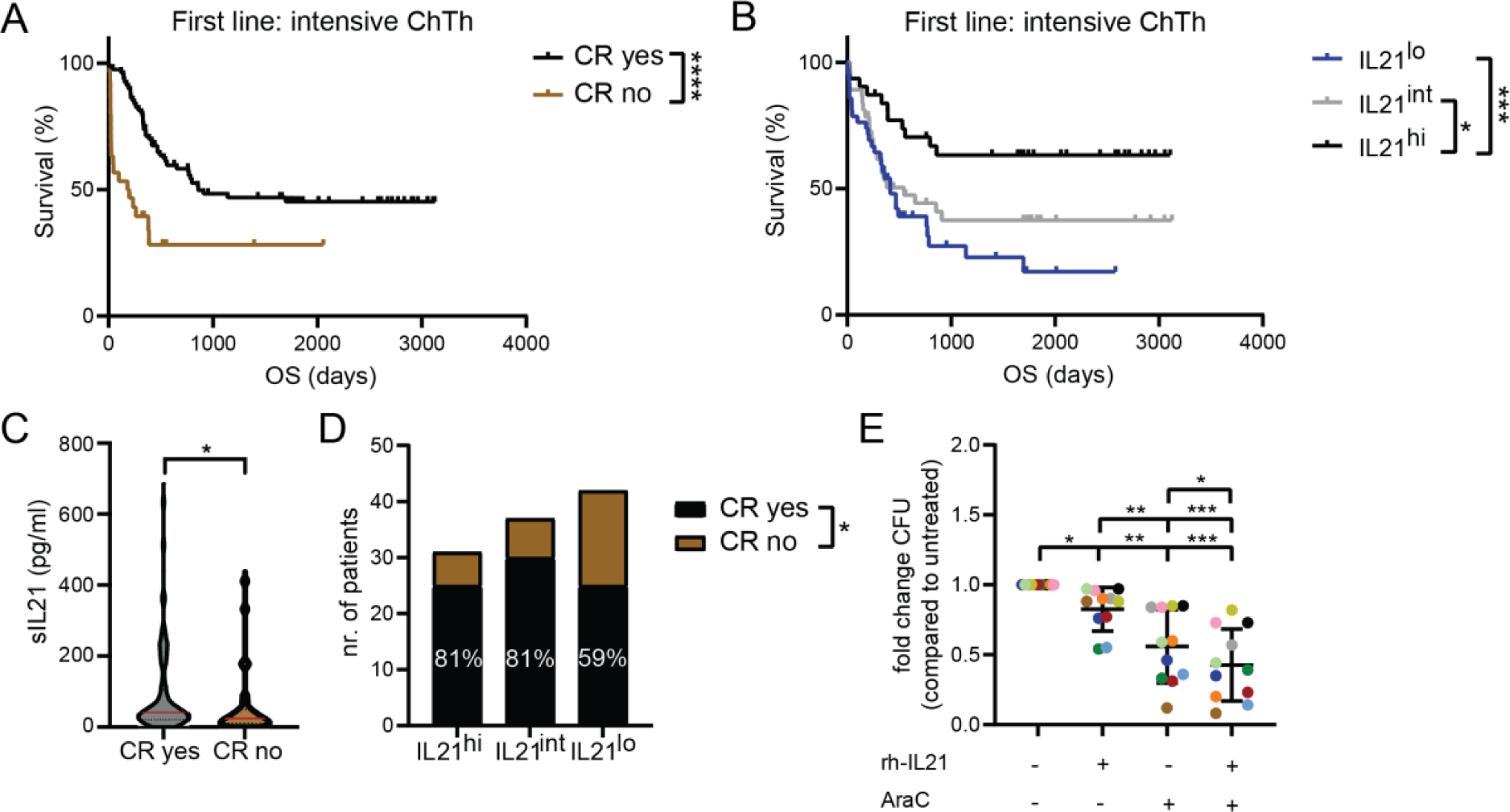
IL21/IL21R signaling promotes the sensitivity of AML LSCs to cytarabine treatment. **(A)** Kaplan-Meier survival curves of AML patients (n = 110) that received intensive chemotherapy (“7+3” regimen) as a first line treatment divided into two groups based on complete remission (CR) achievement. Statistics were determined by log-rank test. **(B)** Kaplan-Meier survival curves of the AML patient cohort that received intensive chemotherapy (n = 110) divided into three groups at the sIL21 threshold of 29 pg/ml and 72 pg/ml. Statistics were determined by log-rank test. **(C)** sIL21 levels in patients that achieved CR (n = 80) versus patients that did not achieve CR (n = 30). Red bars indicate the mean. Statistics were determined by Mann-Whitney test. **(D)** CR rates in patients with high, intermediate and low sIL21 levels. Statistics were determined by Chi-square test. **(E)** LSPCs colonies were enumerated after two weeks of culture in methylcellulose in the presence or absence of 100 pg/ml rh-IL21, 1 nM cytarabine (AraC) or combination. Each dot represents the mean of three technical replicates. Different colors indicate different patients (n = 11). Data are shown as mean ± SD. Statistics were determined by one-way Anova followed by Tukey’s multiple comparisons test. *, P < 0.05; **, P < 0.01; ***, P < 0.001; ****, P < 0.0001. Abbreviations: ChTh, chemotherapy; OS, overall survival; CR, complete remission; CFU, colony-forming units, AraC, cytosine arabinoside (or cytarabine). See also Figure S7.

To functionally investigate whether IL21 render LSPCs more susceptible to chemotherapeutic treatment in vitro, we incubated FACS-sorted LSPCs from different newly diagnosed AML patients with cytarabine and rh-IL21 for 24 hours prior to culturing them in methylcellulose. We observed that the combination of IL21 and cytarabine induced a stronger reduction in LSPCs colony-forming capacity when compared with IL21 or cytarabine single treatments (**Figure 6E**).

These findings support the hypothesis that IL21/IL21R signaling contributes to better response to chemotherapy and is associated with higher CR rates in AML patients.

## Discussion

Quiescent, therapy-resistant LSCs are the major cause of relapse after initially successful chemotherapy in AML^48–50^. Therefore, novel methods effectively eradicating LSCs are an unmet medical need.

Self-renewal is a key feature of both normal and malignant stem cells which allows to maintain and expand the stem cell pool through, respectively, asymmetric division or symmetric renewal^51,52^. Stem cell-related gene signatures are considered as poor prognostic markers for response to therapy and OS in AML^53–55^. Similarly, high frequencies and numbers of LSCs at diagnosis predict therapy resistance and negatively correlate with outcome^56,57^. Mechanistically, these stem cell-related signatures are frequently sustained at the expense of differentiation-promoting events, resulting in a block of differentiation and senescence of AML cells^58^. All-trans retinoic acid is the first approved drug that induces differentiation of blasts in a specific subtype of AML, namely PML-RARα-positive acute promyelocytic leukemia^59^. In addition, inhibition of isocitrate dehydrogenase (IDH) 1 and 2 in AML patients carrying mutations in IDH1 or 2 induces differentiation of AML cells^60^. Recently, several other signaling cascades have been identified which may promote the differentiation of AML cells^61–63^. In this study, we demonstrate that activation of IL21/IL21R signaling favors asymmetric cell division over symmetric renewal in LSCs. This change in division pattern resulted in a reduced pool of AML LSCs through promotion of differentiation, as indicated by increased expression of the differentiation marker CD11b and increased intracellular ROS content, as well as a reduction of LSCs frequency in secondary transplantation experiments. This is in line with recent findings that suggested a correlation between metabolic state, cellular ROS and the differentiation status of HSCs and LSCs. Several studies showed that LSCs have increased mitochondrial mass compared to their healthy counterparts and rely on OXPHOS for energy production^64–66^. Nevertheless, through high activity of ROS-removing pathways, they can maintain very low ROS levels, a characteristic of self-renewing quiescent stem cells, whereas elevated ROS levels can push cells out of quiescence and promote differentiation^64,67,68^.

Many signaling pathways have been shown to promote stemness or differentiation of HSCs and LSCs such as NF-kB, WNT and p38-MAPK signaling^69–72^. ROS modulate some of these molecules with reported function in stem cell maintenance and differentiation such as p38-MAPK. Activation of the p38-MAPK pathway by ROS is crucial in limiting the lifespan and functionality of HSCs as demonstrated in serial transplantation experiments^73–75^. Similarly, treatment of HSCs with a p38-MAPK inhibitor substantially improved the lifespan of HSCs^76^. In complementary experiments, HSC-specific phosphorylation of p38-MAPK resulted in HSC activation and subsequent loss in self-renewal and HSC maintenance^77^. Furthermore, ROS-low HSCs were shown to have higher self-renewal potential, whereas ROS-high HSCs exhausted faster in serial transplantation assays than their ROS-low counterparts due to increased activation of the p38-MAPK^78^. In myelodysplastic syndromes (MDS), inhibition of cytokine-mediated p38-MAPK signaling increased the clonogenic potential of primary human MDS progenitor cells, suggesting that the p38-MAPK pathway may negatively regulate differentiation^79^. Cytokines together with ROS have been identified as major inducers of p38-MAPK signaling-mediated differentiation in many different cell types^80^. In our study, we show that the cytokine IL21 promotes p38-MAPK signaling in AML LSCs resulting in reduced stemness and increased ROS levels in LSCs. AML is a highly complex and heterogeneous disease that arise from the stepwise acquisition of somatic mutations, including chromosomal aberrations and single nucleotide variants^5^. To develop an IL21/IL21R signaling-stimulating approach towards clinical application in AML, it would therefore be necessary to unravel which AML subtypes preferentially respond to IL21 treatment. Our findings are mostly based on the functional experiments in MLL-driven AML mouse models and a limited number of primary human AML samples with diverse cytogenetic and molecular aberrations. Therefore, further investigations on the role of IL21/IL21R signaling in specific AML subtypes are warranted.

IL21 is secreted by multiple immune cell types and is variously involved in immune responses^17,18,25^. Therefore, the induction of IL21/IL21R signaling on LSCs depends on IL21 provided by cells of the immune microenvironment. In other hematological tumor entities, niche-derived IL21 has either growth-promoting or pro-apoptotic effects^36,37,39^. In our study, analysis of IL21 mRNA and protein levels of immune cells and AML cells combined with adoptive transfer experiments of CD4^+^ T cells identified CD4^+^ T cells as the primary source of IL21 and CD4^+^ T cell-derived IL21 as a negative regulator of stemness in AML. However, the mechanisms how IL21 secretion and release is induced in patients with leukemia are currently unknown.

Furthermore, we found that levels of IL21 in the serum in AML patients are increased compared to healthy controls and serve as an independent positive prognostic marker for OS. Serum IL21 could therefore be used clinically as a surrogate biomarker to address the stemness signature of a patient’s AML blasts and to predict outcome.

Resistance to standard induction chemotherapy is one of the key features of LSCs and is accountable for relapse^81^. However, strategies to overcome therapy resistance in AML are lacking. One possibility to increase the chemotherapy sensitivity in AML could be the promotion of differentiation-promoting signaling cascades in LSCs. In this study, we document that high dose chemotherapy was more effective in patients with high levels of sIL21 at diagnosis as illustrated by a significantly higher rate of CR and prolonged OS.

In AML, chemoresistant AML cells have lower ROS levels in response to cytarabine^82^. LSCs are protected from chemotherapy-induced cell death due to the activation of master regulators such as NRF2, which are involved in neutralizing cellular ROS and restoring redox balance^83^. However, a recent study showed that standard induction chemotherapy leads to elevation of ROS, but does not efficiently target LSCs, indicating that increased ROS levels alone are not sufficient to compromise LSCs’ viability or function^66^. Because we found that IL21/IL21R signaling in AML LSCs increases ROS and promotes proliferation and asymmetric division, we hypothesized that IL21 could render AML LSCs more susceptible for chemotherapy. Further investigations involving primary AML LSCs in vitro indeed demonstrated that activation of IL21/IL21R signaling renders LSCs more susceptible to cytarabine treatment. However, attempts to evaluate the therapeutic potential of systemic delivery of rm-IL21 to AML mice via adeno-associated viral vectors failed due IL21-associated toxicities (data not shown). In patients with solid tumors, issues with hepatic or gastrointestinal toxicities have led to discontinuation of IL21’s clinical development for systemic administration^84^. Nevertheless, as an increasing number of selective drug delivery and targeted immunotherapy strategies are being currently developed and optimized, a potential exists for translating IL21 into clinical application, for example by mean of bi-or tri-specific antibodies or chimeric antigen receptor (CAR) T cells that target AML LSCs.

In summary, CD4^+^ T cell-derived IL21 reduces stemness and therapy resistance of AML LSCs by inhibition of cytokine-induced p38-MAPK signaling and by promoting asymmetric cell division. Therefore, stimulating the IL21/IL21R signaling pathway may be a novel immunotherapeutic approach that allows the selective elimination of LSCs.

## Acknowledgments

We thank the staff of the FACS lab (Department for BioMedical Research (DBMR), University of Bern, Switzerland) for providing excellent technical assistance. This work was supported by grants from the Swiss Cancer Research foundation (KFS-4389-02-2018) and Swiss National Science Foundation (310030_179394).

## Author contributions

Conceptualization: C.R. Methodology: V.R., I.K., M.K., A.F.O. and C.R.; Investigation: V.R., M.H., L.T., S.H., I.K., R.R., U.B.; Writing – Original Draft: V.R. and C.R.; Writing – Review: all authors; Supervision: C.R.

## RESOURCES TABLE

1 RESOURCES TABLE
| REAGENT or RESOURCE | SOURCE | IDENTIFIER |
| --- | --- | --- |
| **Antibodies** |  |  |
| Anti-mouse Ly-6C/G (Gr1-1)-APC | BioLegend | Cat# 108412<br>RRID: AB_313377 |
| Anti-mouse CD11b-PE-Cy7 | BioLegend | Cat# 101216<br>RRID: AB_312798 |
| Anti-mouse CD11b- PerCP-Cy5.5 | BioLegend | Cat# 101228<br>RRID:AB_893232 |
| Anti-mouse CD19-APC-Cy7 | BioLegend | Cat# 115530<br>RRID:AB_830707 |
| Anti-mouse Ly-6A/E (Sca-1)-PerCP-Cy5.5 | BioLegend | Cat# 122524<br>RRID:AB_893617 |
| Anti-mouse Ly-6A/E (Sca-1)-APC-Cy7 | BioLegend | Cat# 108126<br>RRID: AB_10645327 |
| Anti-mouse CD117 (c-kit)-APC-Cy7 | BioLegend | Cat# 105826<br>RRID:AB_1626278 |
| Anti-mouse CD16/32 (Fcγ)-PE-Cy7 | BioLegend | Cat# 101318<br>RRID:AB_2104156 |
| Anti-mouse CD4-BV650 | BioLegend | Cat# 100555<br>RRID:AB_2562529 |
| Anti-mouse CD8a-Alexa Fluor® 700 | BioLegend | Cat# 100730<br>RRID:AB_493703 |
| Anti-mouse CD19-biotin | BioLegend | Cat# 115503<br>RRID: AB_313638 |
| Anti-mouse CD11b-PE-Cy7 | BioLegend | Cat# 101216<br>RRID: AB_312799 |
| Anti-mouse CD3ε-biotin | BioLegend | Cat# 100304<br>RRID: AB_312669 |
| Anti-mouse Ter119-biotin | BioLegend | Cat# 116203 RRID:<br>AB_313704 |
| Anti-mouse CD34-eFluor® 450 | ThermoFisher Scientific | Cat# 48-0341-82<br>RRID: AB_2043837 |
| Anti-mouse CD117 (c-kit)-BUV395 | BD Biosciences | Cat# 564011<br>RRID:AB_2738541 |
| Anti-human CD34-APC | BioLegend | Cat# 343608<br>RRID: AB_2228972 |
| Anti-human CD38-PE-Cy7 | BioLegend | Cat# 303515<br>RRID: AB_1279235 |
| Anti-human CD2-biotin | Biolegend | Cat# 300204<br>RRID: AB_314028 |
| Anti-human CD14-biotin | BioLegend | Cat# 325624<br>RRID: AB_2074052 |
| Anti-human CD16-biotin | BioLegend | Cat# 302004<br>RRID: AB_314204 |
| Anti-human CD19-biotin | BioLegend | Cat# 302203<br>RRID: AB_314233 |
| Anti-human CD56-biotin | BioLegend | Cat# 318320<br>RRID: AB_893390 |
| Anti-human CD235ab-biotin | BioLegend | Cat# 306618<br>RRID: AB_2565773 |
| Anti-human CD4-APC-Cy7 | BioLegend | Cat# 317417<br>RRID: AB_571946 |
| Anti-human CD8-PerCP-Cy5.5 | BioLegend | Cat# 344710<br>RRID: AB_2044010 |
| Anti-human CD45-V500-C | BD Biosciences | Cat# 647449<br>RRID: AB_2870319 |
| Anti-human CD360 (IL21R) | Miltenyi Biotec | Cat# 130-101-477<br>RRID: AB_2657745 |
| Human IgG1, REA Control Antibody (S) | Miltenyi Biotec | Cat# 130-113-438<br>RRID: AB_2733893 |
| Rabbit anti-IkBα (44D4) mAb (unconjugated) | Cell Signaling Technology | Cat# 4812<br>RRID: AB_10694416 |
| Rabbit anti-plkBα (14D4) mAb (unconjugated) | Cell Signaling Technology | Cat# 2859<br>RRID: AB_561111 |
| Anti-rabbit IgG (H+L), F(ab')<sub>2</sub> Fragment-Alexa Fluor® 647 | Cell Signaling Technology | Cat# 4414<br>RRID: AB_10693544 |
| Goat anti-Numb polyclonal Ab (unconjugated) | Abcam | Cat# ab4147<br>RRID: AB_304320 |
| Mouse anti-Tubulin mAb (unconjugated) | Abcam | Cat# ab7291<br>RRID: AB_2241126 |
| AlexaFluor®568 donkey-anti goat IgG H&L | Abcam | Cat# ab175704<br>RRID: AB_2725786 |
| AlexaFluor®647 rabbit-anti mouse IgG H&L | Abcam | Cat# ab150127 |
| PE anti-p38 MAPK Phospho (Thr180/Tyr182) | BioLegend | Cat#690203<br>RRID: AB_2832849 |
| PE Mouse IgG1, κ Isotype Ctrl Antibody | BioLegend | Cat#400139<br>RRID: AB_493443 |
| Biological samples |  |  |
| BM aspirates from untreated, newly diagnosed AML patients | Department of Hematology and Central Hematology Laboratory, Inselspital, Bern University Hospital and University of Bern | N/A |
| PB from untreated, newly diagnosed AML patients | Department of Hematology and Central Hematology Laboratory, Inselspital, Bern University Hospital and University of Bern | N/A |
| BM from healthy donors<br>(orthopedic patients who underwent Vertebroplasty) | Inselspital, Bern University Hospital and University of Bern | N/A |
| Chemicals, peptides, and recombinant proteins |  |  |
| MethoCult™ H4435 Enriched for human cells | STEMCELL Technologies | Cat# 04435 |
| MethoCult™ M3134 for mouse cells | STEMCELL Technologies | Cat# 03134 |
| Human Methylcellulose Serum-Free Enriched Media | R&D Systems | Cat# HSC005SF |
| Mouse recombinant IL-3 | Miltenyi | Cat#130-099-508 |
| Mouse recombinant IL-6 | Miltenyi | Cat#130-096-682 |
| Mouse recombinant SCF | Miltenyi | Cat#130-101-697 |
| Mouse recombinant Flt3l | Miltenyi | Cat#130-094-038 |
| Human Insulin Actrapid | Novo Nordisk | Cat#8-0201-83-201-3 |
| Human holo-transferrin | Prospec | Cat#PRO-315 |
| StemSpan™ Medium | STEMCELL Technologies | Cat# 09605 |
| StemSpan™ Cytokine Cocktail (CC)-100 | STEMCELL Technologies | Cat# 02690 |
| APC Streptavidin | BioLegend | Cat# 405207 |
| FITC Streptavidin | BioLegend | Cat# 405202 |
| Pacific Blue™ Annexin V | BioLegend | Cat# 640918 |
| V500 Streptavidin | BD Biosciences | Cat# 561419,<br>RRID:AB_10611863 |
| Fixable Viability Dye eFluor® 506 | ThermoFisher Scientific | Cat# 65-0866-14 |
| Fixable Viability Dye eFluor® 450 | ThermoFisher Scientific | Cat# 65-0863-14 |
| Human recombinant IL-21 | Sigma-Aldrich | Cat# SRP3087 |
| Cytosine β-D-arabinofuranoside | Sigma-Aldrich | Cat# C1768 |
| DAPI | Merck | Cat# 10236276001 |
| Dako Wash | Agilent Technologies | Cat# S300685-2 |
| Dako Antibody Diluent | Agilent Technologies | Cat# S080983-2 |
| CellROX™ Deep Red Reagent | ThermoFisher Scientific | Cat# C10422 |
| Tetramethylrhodamine, Methyl Ester, Perchlorate (TMRM™) | ThermoFisher Scientific | Cat# T668 |
| MitoTracker™ Red FM | ThermoFisher Scientific | Cat# M22425 |
| SB203580 p38 MAPK inhibitor | STEMCELL Technologies | Cat#72222 |
| True-Phos™ Perm Buffer | BioLegend | Cat#425401 |
| Critical commercial assays |  |  |
| APC BrdU Flow Kit | BD Biosciences | Cat# 552598<br>RRID: AB_2861367 |
| RNA Easy Micro Kit | Qiagen | Cat# 74004 |
| Quick-RNA Microprep Kit | Zymo Research | Cat# R1051 |
| ELISA MAX™ Deluxe Set Human IL-21 | BioLegend | Cat# 433804 |
| Deposited data |  |  |
| RNA-seq data |  | GEO: GSE241170,<br>GSE241171, GSE241172 |
| Experimental models: Cell lines |  |  |
| THP-1 | ATCC | TIB-202™ |
| Experimental models: Organisms/strains |  |  |
| Mouse: C57BL/6J | Charles River | MGI:3028467 |
| *Il21R<sup>-/-</sup>* (B6.129-Il21rtm1Kopf/J) |  | MGI:5435248 |
| *Il21<sup>-/-</sup>* (B6.129S-Il21tm1Lex/Mmucd) |  | MGI:4843320 |
| *Il21<sup>mCherry</sup>* (B6.Cg-Tg(Il21-mCherry)1Wjl/Mmnc) |  | MGI:5478531 |
| Oligonucleotides |  |  |

| Recombinant DNA |  |  |
| --- | --- | --- |
| Software and algorithms |  |  |
| FlowJo™ software v.10.6 | TreeStar |  |
| GraphPad Prism® software v9.0 | GraphPad |  |
| ELDA Software | Hu and Smyth et al., 2009 | http://bioinf.wehi.edu.au/software/elda/ |
| X-tyle software | Camp and Dolled-Filhart, 2004 | https://medicine.yale.edu/lab/rimm/research/software/ |
| fastqc v. 0.11.9 |  | https://www.bioinformatics.babraham.ac.uk/projects/fastqc/ |
| RSeQC v. 4.0.0 | Wang et al., 2012 |  |
| HiSat2 v. 2.2.1 | Kim et al., 2015 |  |
| FeatureCounts v. 2.0.1 | Liao et al., 2014 |  |
| DESeq2 v.1.32.0 | Love et al., 2014 |  |
| TopGo v. 2.44.0 | Alexa and Rahnenfuhrer, 2021 |  |
| ClusterProfiler v. 4.0.2 | Yu et al., 2012 |  |
| R 4.1.0 |  | https://www.R-project.org/ |
| INSPIRE® software |  | https://www.merckmillipore.com/ |
| IDEAS® software |  | https://www.merckmillipore.com/ |
| Kaluza analysis software | Beckman Coulter Life Sciences |  |
| Other |  |  |
| msigdb_HALLMARK_OXIDATIVE_PHOSPHORYLATION |  | [gsea-msigdb.org](https://www.gsea-msigdb.org) |
| Somerville_UP | Somerville and Cleary, 2006 |  |
| Somerville_DOWN | Somerville and Cleary, 2006 |  |
| STEMNESS_UP |  | [gsea-msigdb.org](https://www.gsea-msigdb.org) |
| STEMNESS_DOWN |  | [gsea-msigdb.org](https://www.gsea-msigdb.org) |
| GAL_LEUKEMIC_STEM_CELL_DOWN |  | [gsea-msigdb.org](https://www.gsea-msigdb.org) |
| BROWN_MYELOID_CELL_DEVELOPMENT_DN |  | [gsea-msigdb.org](https://www.gsea-msigdb.org) |
| BROWN_MYELOID_CELL_DEVELOPMENT<br>_UP |  | gsea-msigdb.org |
| msigdb_HALLMARK_TNFA_SIGNALING<br>_VIA_NFKB |  | gsea-msigdb.org |
| HSC Differentiation and Lineage-specific<br>markers |  | pathcards.genecards.org |
| Wnt Signaling (beta-Catenin) |  | pathcards.genecards.org |
| MAPK-Erk Signaling |  | pathcards.genecards.org |
| POSITIVE REGULATIN OF CELL CYCLE |  | gsea-msigdb.org |
| Respiratory electron transport_ATP synthesis |  | pathcards.genecards.org |

## RESOURCE AVAILABILITY

### Lead contact

Further information and requests for resources and reagents should be directed to and will be fulfilled by the lead contact, Carsten Riether.

### Materials availability

All unique reagents generated in this study are available from the Lead Contact without restriction.

### Data and code availability

All RNA-seq data compiled for this study is made publicly available on the Gene Expression Omnibus (GEO) website (https://www.ncbi.nlm.nih.gov/geo/) under the accession number GSE241170, GSE241171 and GSE241172. This study does not include the development of new code.

## EXPERIMENTAL MODEL AND SUBJECT DETAILS

### Mice

C57BL/6J (BL/6) mice and NOD SCID gamma (NSG) mice were purchased from Charles River Laboratories (Sulzfeld, Germany). *Il*21^-/-^ mice on BL/6J background^85^ were kindly provided by Prof. Daniel Pinschewer (Department of Biomedicine, University of Basel). *Il21R*^-/-^ mice on BL/6J background^23^ and IL21^mCherry^ reporter mice^43^ were kindly provided by Prof. Manfred Kopf (Molecular Health Sciences, ETH Zurich). Experiments were performed with age-(6-8 weeks) and sex-matched animals of both genders and mice were randomly assigned to different treatment groups. Mice were housed under specific pathogen-free conditions in individually ventilated cages with food and water ad libitum and were regularly monitored for pathogens. Animal experiments were approved by the local experimental animal committee of the Canton of Bern and performed according to Swiss laws for animal protection (BE75/17, BE78/17, BE56/2020, BE59/2020, BE58/2021 and BE30/2021).

### Patient samples

Blood samples, BM aspirates and serum from untreated, newly diagnosed AML patients at the Department of Hematology and Central Hematology Laboratory, Inselspital, Bern University Hospital and University of Bern, Switzerland, were obtained after written informed consent. BM from healthy donors was collected from orthopedic patients who underwent Vertebroplasty. Patient characteristics are listed in Table S1 and Table S2. Analysis of blood, BM and serum samples was approved by the local ethical committee of the Canton of Bern, Switzerland (KEK 122/14 and 2019-01627).

### Cell lines

THP-1 cells were purchased from ATCC, cultured in RPMI 1640 medium supplemented with 10% fetal calf serum (FCS), 100 U/ml penicillin, 100 µg/ml streptomycin and maintained in a humidified incubator at 37℃ and 5% CO_2_.

## METHOD DETAILS

### Antibodies for flow cytometry and cell sorting

#### Human

APC anti-CD34 (clone 561, 1:80), PE-Cy7 anti-CD38 (clone HIT2, 1:50), APC-Cy7 anti-CD4 (clone RPA-T4, 1:80), PerCP-Cy5.5 anti-CD8 (clone HIT8a, 1:100), Pacific Blue Annexin V (1:50) were purchased from BioLegend. V500 anti-CD45 (clone 2D1, 1:50) was purchased by BD Biosciences. PE anti-IL21R (clone REA233, 1:10) and PE anti-CD132 (clone REA313, 1:10), PE REA Control (S) human IgG1 isotype were purchased from Miltenyi. Differentiated cells were excluded by using biotin-conjugated antibodies against CD2 (clone RPA-2.10), CD14 (clone HCD14), CD16 (clone 3G8), CD19 (clone HIB19), CD56 (clone HCD56), CD235a (clone HIR2) (all 1:100; BioLegend), followed by staining with FITC-conjugated streptavidin (1:3000; BioLegend).

#### Mouse

APC anti-Ly-6C/G (Gr1-1) (clone RB6-8C5, 1:200), PE-Cy7 anti-CD11b (clone M1/70, 1:200), APC-Cy7 anti-CD19 (clone 6D5, 1:300), PerCP-Cy5.5 anti-Sca-1 (clone D7, 1:600), APC-Cy7 anti-CD117 (c-kit) (clone 2B8, 1:300), PE-Cy7 anti-CD16/32 (Fcγ) (clone 93, 1:400), BV650 anti-CD4 (clone RM4-5, 1:600), AlexaFluor700 anti-CD8 (clone 53-6.7, 1:800) were purchased from BioLegend. e506 Fixable Viability Dye (1:1000), eFluor450 anti-CD34 (clone RAM34, 1:100) were purchased from ThermoFisher Scientific. Lineage positive were excluded by using biotin-conjugated antibodies against CD19 (clone 6D5), CD3e (clone 145-2C11), Ly-6C/G (Gr1-1) (clone RB6-8C5), Ter119 (clone Ter-119) (all 1:300; BioLegend), followed by staining with V500-streptavidin (1:1000; BD Biosciences) or APC-streptavidin (1:3000; BioLegend).

Samples were acquired on a BD LSR Fortessa and sorting procedures were conducted using a BD FACS Aria III (both BD Biosciences). Data were analyzed using FlowJo software v.10.6 (TreeStar).

### IL21 determination in human serum and mouse BM

Human IL21 protein levels in serum samples from newly diagnosed, untreated AML patients were determined by enzyme-linked immunosorbent assay (ELISA) using an IL21 Human ELISA kit (Biolegend), according to the manufacturer’s instructions.

To obtain murine BM supernatant, bones were flushed in 400 µl of phosphate-buffered saline (PBS) solution and supernatant was collected after pelleting the cells. Mouse IL21 protein levels in BM supernatant were determined using an IL21 Mouse ELISA kit (ThermoFisher Scientific).

### Colony-forming Assay

#### Human

3×10^3^ FACS-purified CD45^dim^SSC^lo^lin^−^CD34^+^ AML stem and progenitor cells from BM and PB of AML patients were cultured overnight in 96-well V-bottom plates (Corning) in Stem Span Medium (STEMCELL Technologies) supplemented with 100X Stem Span Cytokine Cocktail (STEMCELL Technologies), in the presence or absence of 100 pg/ml recombinant human (rh)-IL21 (Sigma-Aldrich). The next day, cells were plated into semi-solid methylcellulose (MethoCult™ H4435 Enriched, STEMCELL Technologies or Enriched Human Methylcellulose, R&D Systems), with further addition of 100 pg/ml rh-IL21. For the second plating, total cells were collected from the methylcellulose and counted, followed by replating of 10-fold higher cell number into methylcellulose, without any treatment. Colonies number (≥ 30 cells/colony) was assessed by inverted light microscopy after 2 weeks for each round of plating.

#### Mouse

For assessment of colony forming units, 5×10^4^ total bone marrow cells from AML mice were plated into semi-solid methylcellulose. Number of MLL-AF9-GFP^+^ or MLL-ENL-YFP^+^ colonies (≥ 30 cells/colony) was assessed after 7 days by fluorescence microscopy.

For testing the effect of recombinant mouse IL21 on LSCs colony-forming capacity, 10^3^ FACS-purified GFP^+^lin^-^Sca-1^-^c-kit^high^CD34^+^Fcγ^+^ GMPs from AML mice were cultured overnight in 96-well V-bottom plates (Corning) in RPMI 1640 medium supplemented with 10% FCS, 1% L-glutamine, 1% penicillin/streptomycin, 100 ng/ml recombinant mouse (rm)-SCF (Miltenyi) and 20 ng/ml rm-TPO (BioLegend), in the presence or absence of 300 pg/ml rm-IL21 (R&D Systems). The next day, cells were plated in semi-solid methylcellulose. Number of colonies was assessed by inverted light microscopy after 7 days.

In both assays, the semi-solid methylcellulose used was MethoCult M3134 medium (STEMCELL Technologies), supplemented with 15% FCS, 20% BIT (50 mg/ml BSA in IMDM, 1.44 U/ml rh-insulin [Actrapid; Novo Nordisk], and 250 ng/ml human holo transferrin [Prospec]), 100 µM 2-mercaptoethanol, 100 U/ml penicillin, 100 µg/ml streptomycin, 2 mM L-glutamine, and 50 ng/ml rm-SCF, 10 ng/ml rm-IL-3, 10 ng/ml rm-IL-6, and 50 ng/ml rm-Flt3-ligand (all Miltenyi).

### Short-term LSPCs liquid culture

FACS-purified CD45^dim^SSC^lo^lin^-^CD34^+^ AML stem and progenitor cells from BM and PB of AML patients were cultured for 72 hours in 96-well V-bottom plates (Corning) in StemSpan Medium (STEMCELL Technologies) supplemented with 100X StemSpan Cytokine Cocktail (STEMCELL Technologies), in the presence or absence of 100 pg/ml rh-IL21 (Sigma-Aldrich) and/or 1 nM cytarabine (Cytosine β-D-arabinofuranoside, Sigma-Aldrich). Cells were counted after 72 h to determine cell growth and stained with AnnexinV to assess viability.

### AML models

MLL-AF9 AML was induced by transducing FACS-purified LSKs with the GFP-MLL-AF9 retroviral construct^40^ by spin infection on two consecutive days. 5×10^4^ cells were injected into the tail vein of non-irradiated syngeneic recipients.

MLL-ENL AML was induced by retroviral transduction of FACS-purified LSKs with the YFP-MLL-ENL oncogene^86^, with spin infection on two consecutive days. 2.5×10^4^ cells were injected into the tail vein of sublethally irradiated (4.5 Gy) syngeneic recipients.

### LSPCs analysis

#### Human

AML stem and progenitor cells from BM and PB primary samples for phenotypical analysis and/or FACS-purification were defined as CD45^dim^SSC^lo^lin^-^CD34^+^ according to several publications^87–89^.

#### Mouse

L-GMPs in BM and spleen of AML mice were for phenotypical analysis and/or FACS-purification were defined as GFP/YFP^+^lin^-^Sca-1^-^c-kit^high^CD34^+^Fcγ^+^ cells according to Somervaille and Cleary, 2006^40^.

### Numb staining and ImageStream analysis

FACS-sorted L-GMPs were fixed by incubation in 4% paraformaldehyde (PFA), followed by permeabilization with 1X wash buffer (Dako Wash, Agilent Technologies) and blocking with 10% normal rabbit serum and donkey serum (Invitrogen) in Dako Wash. Numb and α-tubulin staining was performed overnight at 4 °C (with goat anti-Numb polyclonal ab, 1:20; and mouse anti-tubulin mAb, 1:400; both Abcam) in diluent (Dako antibody diluent, Agilent Technologies). Cells were then incubated with the secondary antibody (AlexaFluor568-conjugated donkey-anti goat ab, 1:400, and AlexaFluor647-conjugated rabbit-anti mouse ab, 1:2000; both Abcam) for 1 hour at room temperature. DAPI (Roche) was used to stain for DNA. Samples were acquired using an ImageStream®xMkII imaging flow cytometer (Merck) and dividing cells were analyzed using INSPIRE^®^ and IDEAS^®^ Software. A difference in Numb staining of at least 1.8 fold was defined as asymmetric cell division as described by Zimdahl et al., 2014^90^.

### brdU staining

1 mg brdU solution (BD Biosciences) was injected intraperitoneally 48 hours prior to sacrificing and analyzing the mice. brdU staining was performed using the brdU APC kit (BD Biosciences), according to the manufacturer’s instructions. Briefly, after cell surface markers staining, whole BM cells were fixed and permeabilized by incubation in BD Cytofix/Cytoperm. Next, cells were incubated with 300 µg/mL DNase for 1 hour at 37°C to expose incorporated brdU and then stained with an APC anti-brdU antibody (1:50, BD Biosciences). 1X BD Perm/Wash in ddH_2_O was used as a staining and washing buffer. Samples were acquired on an LSRII (BD Biosciences) flow cytometer.

### NF-kB staining

Whole BM cells were fixed by incubation in 4% PFA for 15 minutes, followed by permeabilization in ice-cold 90% methanol in 1X PBS. The following antibodies were used for intracellular staining respectively of IKBα and phosphoIKBα: rabbit mAb anti-IKBα (44D4) (1:100, Cell Signaling) and rabbit mAb anti-phospho-IKBα (Ser32) (14D4) (1:100, Cell Signaling), followed by staining with an AlexaFluor647-conjugated anti-rabbit IgG F(ab’)2 Fragment (1:800, Cell Signaling). 0.5% BSA in 1X PBS was used as a staining and washing buffer. Samples were acquired on an LSRII (BD Biosciences) flow cytometer.

### p38-MAPK staining

Intracellular p38-MAPK staining was performed for FACS-sorted L-GMPs from BM of BL/6, *Il21^-/-^*, *Il21R^-/-^* and *Il21R^+/-^* AML mice and for THP-1 AML cells, which were first treated with 1 ng/ml rh-IL21 for 72 hours or left untreated. After staining with eFluor450 Fixable Viability Dye (1:1000; ThermoFisher Scientific), cells were fixed with Cytofix/Cytoperm (BD Bioscience) as per manufacturer protocol. Subsequently cells were washed with PBS and permeabilized with True-Phos™ Perm Buffer (BioLegend) according to the manufacturer’s protocol. Cells were washed with PBS and intracellularly stained with PE anti-phospho-p38-MAPK (Thr180/Tyr182; 1:20; BioLegend) ab or PE mouse isotype control IgG1,κ (1:20; BioLegend) for 30 minutes at room temperature. Cells were acquired on an LSRII (BD Biosciences) flow cytometer and analyzed by Kaluza Analysis software.

### ROS and mitochondrial dyes staining

Staining with CellROX^TM^, MitoTracker^TM^ and TMRM^TM^ was performed on whole BM cells, after cell surface markers staining. For staining, cells were incubated for 30 min at 37°C with, respectively, 5 μM CellROX^TM^, 25 nm MitoTracker^TM^, 10 nm TMRM^TM^ in RPMI 1640 medium. 50 μM of verapamil hydrochloride was used to inhibit dyes efflux from mitochondria. Cells were washed three times with PBS to remove the excess of dyes and were acquired on an LSRII (BD Biosciences) flow cytometer.

### Cell culture with p38 MAPK inhibitor

#### THP-1 cell line

10^5^ human THP-1 AML cells were pretreated for 30 minutes with vehicle or 10 nm/ml of the p38 MAPK inhibitor SB203580 and were then cultured for 72 h in the presence or absence of 1 ng/ml rh-IL21 in technical triplicates. After 72 h of culture, cell numbers were assessed by Trypan-blue exclusion and intracellular ROS and p38-MAPK phosphorylation by FACS.

#### Murine L-GMPs

10^3^ FACS-purified GFP^+^lin^-^Sca-1^-^c-kit^high^CD34^+^Fcγ^+^ GMPs from AML mice were pretreated for 30 minutes with vehicle or 10 nm/ml of the p38 MAPK inhibitor SB203580 and were then cultured overnight in 96-well V-bottom plates (Corning) in RPMI 1640 medium supplemented with 10% FCS, 1% L-glutamine, 1% penicillin/streptomycin, 100 ng/ml recombinant mouse (rm)-SCF (Miltenyi) and 20 ng/ml rm-TPO (BioLegend), in the presence or absence of 300 pg/ml rm-IL21 (R&D Systems). The next day, cells were plated in semi-solid methylcellulose (described above). Number of colonies was assessed by inverted light microscopy after 7 days.

### Quantitative Reverse Transcription PCR analysis of gene expression

For quantitative Reverse Transcription PCR (qRT-PCR), total RNA was extracted from FACS-sorted cell populations using the Quick-RNA MiniPrep kit (Zymo Research) according to the manufacturer’s instructions. Total RNA was reverse-transcribed using 2.5×10-4 U/μl hexanucleotide mix (Roche), 0.4mM deoxynucleotide mix (Sigma-Aldrich), 1.25 U/μl RNAsin (Promega) and 4 U/μl reverse transcriptase (Promega). 2 µl of cDNA were used for Real-Time PCR with self-designed primers and SYBR green reaction (Roche). qRT-PCR reactions were performed in duplicates including non-template controls on a QuantStudio 3 Real-Time PCR system (Applied Biosystems). Expression levels of analyzed genes relative to a reference gene (ACTB or Actb) were calculated using the comparative Ct method (also referred to as the 2^-ΔΔCt^ method)^91^. The following primer pairs were used to determine mRNA expression of respective genes:

*Il21*, FW: CACATAGCTAAATGCCCTTCC, RV: CCTCAGGAATCTTCGGGTC;

*Actb*, FW: AGATGACCCAGATCATGTTTGAG, RV: GTACGACCAGAGGCATACAG;

*IL21*, FW: TTATGTGAATGACTTGGTCCCT, RV: CTGTATTTGCTGACTTTAGTTGGG;

*IL21R*, FW: TCATCTTTCAGACCCAGTCAG, RV: CATATCTTCTTCCATAGCCTCCAC;

*ACTB*, FW: GCACCACACCTTCTACAATGAG, RV: GGTCTCAAACATGATCTGGGTC.

### High-throughput transcriptome analysis using next generation RNA sequencing

Total RNA was extracted from FACS-sorted L-GMPs from BL/6 and *Il21*^-/-^ AML mice using the RNeasy Micro Kit (Qiagen). RNA purity was checked using the NanoPhotometer® spectrophotometer (Implen). RNA integrity and quantity were assessed using the RNA Nano 6000 Assay Kit of the Bioanalyzer 2100 system (Agilent Technologies). A total amount of 1 μg RNA per sample was used as input material for the libraries preparations. Sequencing libraries were generated using NEBNext® UltraTM RNA Library Prep Kit for Illumina® (New England Biolabs Inc.) and index codes were added to attribute sequences to each sample. Library quality was assessed on the Agilent Bioanalyzer 2100 system (Agilent Technologies). The clustering of the index-coded samples was performed on a cBot Cluster Generation System using PE Cluster Kit cBot-HS (Illumina). After cluster generation, the libraries were sequenced on a Nova 6000 Illumina platform and paired-end reads were generated.

### RNA-seq analysis and Gene Set Enrichment Analysis

The quality of the RNA-seq data was assessed using fastqc v. 0.11.9 (http://www.bioinformatics.babraham.ac.uk/projects/fastqc/) and RSeQC v. 4.0.0^92^. The reads were mapped to the GRCm38 reference genome using HiSat2 v. 2.2.1^93^. FeatureCounts v. 2.0.1^94^ were used to count the number of reads overlapping with each gene as specified in the genome annotation (Ensembl build 100 and Homo_sapiens.GRCh38.104).

The R Bioconductor package DESeq2 v1.32.0^95^ was used to test for differential gene expression between the experimental groups. TopGo v2.44.0^96^ was used to identify gene ontology terms containing unusually many differentially expressed genes. An interactive Shiny application was set up to facilitate the exploration and visualization of the RNA-seq results.

Gene set enrichment analysis (GSEA) was run with ClusterProfiler v4.0.2^97^, using gene sets from the Broad Institute’s Molecular Signatures Database (MSigDB Hallmarks collection, available at gsea-msigdb.org), Pathcards database (available at https://pathcards.genecards.org/) and KEGG database (available at https://www.genome.jp/kegg/pathway.html). Further visualization was performed using R version 4.1.0.

## QUANTIFICATION AND STATISTICAL ANALYSIS

All flow cytometry, in vitro and in vivo data were analyzed and plotted using GraphPad Prism^®^ software v9.0 (GraphPad). Bars and error bars indicate means, standard errors of mean and standard deviations of the indicated number of independent biological replicates. Two-tailed Student’s *t* test, Mann-Whitney test, Pearson r test, one-way-ANOVA followed by Tukey’s post-test and two-way ANOVA followed by Sidak’s post-test were used as indicated in the figures legends. Significance of differences in Kaplan-Meier survival curves was determined using the log-rank test (two-tailed). LSC frequencies with 95% confidence intervals (CI) were estimated with ELDA software (http://bioinf.wehi.edu.au/software/elda/) and significant differences in LSC frequency were calculated by χ2 test in limiting dilution assay^41^.

P<0.05 was considered significant. Details on the quantification, normalization and statistical tests used in every experiment can be found in the corresponding figure legend. n represents the number of independent replicates in each experiment.

## Abbreviations

LSC: leukemia stem cell
AML: acute myeloid leukemia
IL: interleukin
ROS: reactive oxygen species
HSC: hematopoietic stem cell
FLT3: fms-related tyrosine kinase 3 gene
CR: complete remission
BM: bone marrow
EC: endothelial cell
MSCs: mesenchymal stromal cell
JAK: Janus tyrosine kinases
STAT: signal transducers and activators of transcription
PI3K: phosphoinositol 3-kinase
MAPK: mitogen-activated protein kinase
NK: natural killer
L-GMP: leukemic granulocyte-macrophage progenitor
OS: overall survival
LSKs: lineage^-^Sca-1^+^c-kit^+^ cells
lin: lineage
ELDA: extreme limiting dilution analysis
GO: gene ontology
GSEA: gene set enrichment analysis
sIL21: serum IL21
LSPC: leukemic stem and progenitor cell
HSPC: hematopoietic stem and progenitor cell
IDH: isocitrate dehydrogenase
OXPHOS: oxidative phosphorylation.

