## Supplemental information for "CD4^+^ T cell-derived IL21 regulates stem cell fate in acute myeloid leukemia by activation of p38-MAPK signaling"

#### Figure S1

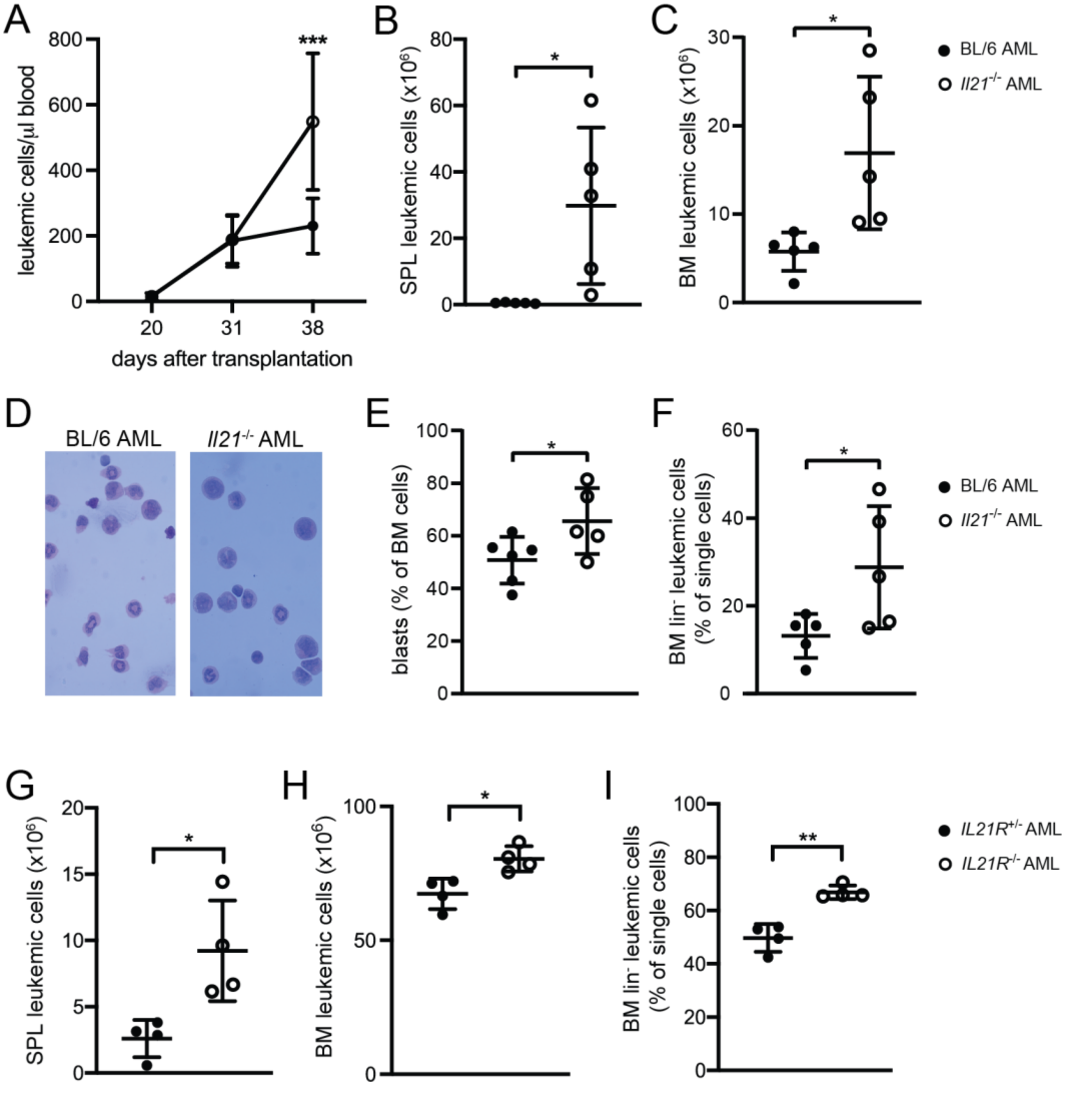


**Figure S1. An IL21-deficient microenvironment, as well as IL21R deficiency on leukemia-initiating cells result in increased AML burden and accumulation of primitive leukemic cells in BM of mice.** **(A)** Number of MLL-AF9-GFP$^+$ leukemic cells on days 20, 31 and 38 in the blood of BL/6 and *Il21*$^{-/-}$ AML mice (n = 5 mice/group). Data are displayed as mean ± SD. Statistics were determined by two-way ANOVA followed by Sidak's multiple comparisons test. **(B, C)** MLL-AF9-GFP$^+$Gr1$^+$Cd11b$^+$ cells in the spleen **(B)**

and in the BM **(C)** of BL/6 and *Il21*$^{-/-}$ AML mice (n = 5 mice/group). Data are displayed as mean ± SD. Statistics were determined by Student's t test. **(D)** Representative H&E-stained cytospin preparations of BM, **(E)** quantification of blasts percentage by microscopic evaluation of cell morphology and **(F)** percentage of lineage negative MLL-AF9-GFP$^{+}$ leukemic cells in BM of BL/6 and *Il21*$^{-/-}$ AML mice (n = 5 mice/group). **(A – F)** One representative of four independent experiments is shown. **(G, H)** MLL-AF9-GFP$^{+}$Gr1$^{+}$Cd11b$^{+}$ cells in the spleen **(G)** and in the BM **(H)** and **(I)** percentage of lineage negative MLL-AF9-GFP$^{+}$ leukemic cells in BM of *Il21R*$^{+/-}$ and *Il21R*$^{-/-}$ AML mice (n = 4 mice/group). **(G – I)** One representative of two independent experiments is shown. Data are displayed as mean ± SD. Statistics were determined by Student's t test.

*, P < 0.05; **, P < 0.01.

Abbreviations: SPL, spleen: BM, bone marrow; Lin, lineage.

### Figure S2

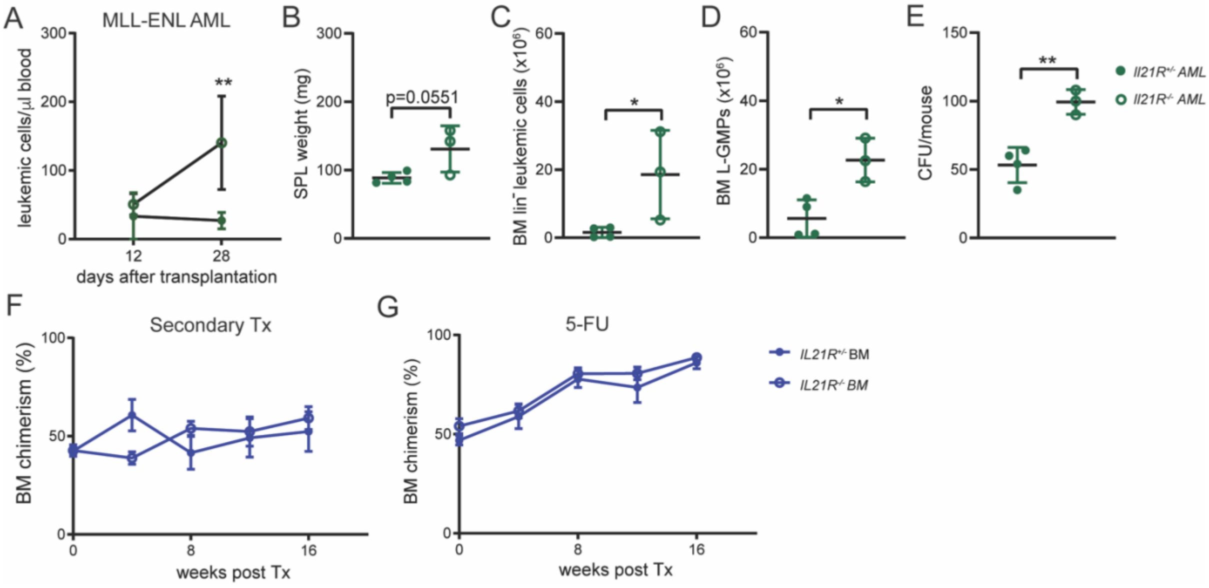


**Figure S2. IL21R deficiency on leukemia-initiating cells results in faster disease development and accumulation of primitive leukemic cells in an MLL-ENL-driven AML model. IL21R deficiency on normal hematopoietic stem cell does not affect their repopulating capacity in steady-state and stress-induced hematopoiesis.**

**(A - D)** $2.5x10^4$ MLL-ENL-YFP-transduced LSKs from the BM of *Il21R*$^{-/-}$ and *Il21R*$^{+/-}$ mice were injected intravenously into sublethally-irradiated (4.5 Gy) *Il21R*$^{+/-}$ recipients (*Il21R*$^{-/-}$ AML and *Il21R*$^{+/-}$ AML, respectively). Mice were sacrificed 30 days after leukemia transplantation and BM and spleen were analyzed (n = 3-4 mice/group). One representative of two independent experiments is shown. **(A)** Number of MLL-ENL-YFP$^+$ leukemic cells on days 12 and 28 in the blood of *Il21R*$^{-/-}$ and *Il21R*$^{+/-}$ AML mice. Data are displayed as mean ± SD. Statistics were determined by two-way ANOVA followed by Sidak's multiple comparisons test. **(B)** Spleen size, **(C)** number of lineage negative MLL-ENL-YFP$^+$ leukemic cells and **(D)** number of L-GMPs in BM of *Il21R*$^{-/-}$ AML and *Il21R*$^{+/-}$ AML mice. Data are displayed as mean ± SD. Statistics were determined by Student's *t* test. **(E)** Colony forming units per mouse. $5x10^4$ BM cells were plated into methylcellulose and YFP$^+$ colonies were enumerated seven days later by inverted fluorescence microscopy. Data are displayed as mean ± SD. Statistics were determined by Student's *t* test. **(F)** BM reconstitution after transplantation of *Il21R*$^{-/-}$ and *Il21R*$^{+/-}$ donor cells into lethally irradiated (2 x 6.5 Gy) congenic secondary recipients. BM chimerism measured at week 4, 8, 12 and 16 post transplantation. Data are displayed as mean ± SEM. Statistics were determined by two-way ANOVA followed by Sidak's multiple comparisons test. **(G)** BM reconstitution after 5-FU treatment followed by transplantation of *Il21R*$^{-/-}$ and *Il21R*$^{+/-}$ donor cells into lethally irradiated (2 x 6.5 Gy) congenic secondary recipients. BM chimerism measured at week 4, 8, 12 and 16 post transplantation. Data are displayed as mean ± SEM. Statistics were determined by two-way ANOVA followed by Sidak's multiple comparisons test.

*, $P < 0.05$; **, $P < 0.01$.

Abbreviations: SPL, spleen; L-GMPs, leukemic granulocyte-macrophage progenitors; CFU, colony-forming units; Tx, transplantation; 5-FU, 5-fluorouracil.

### Figure S3

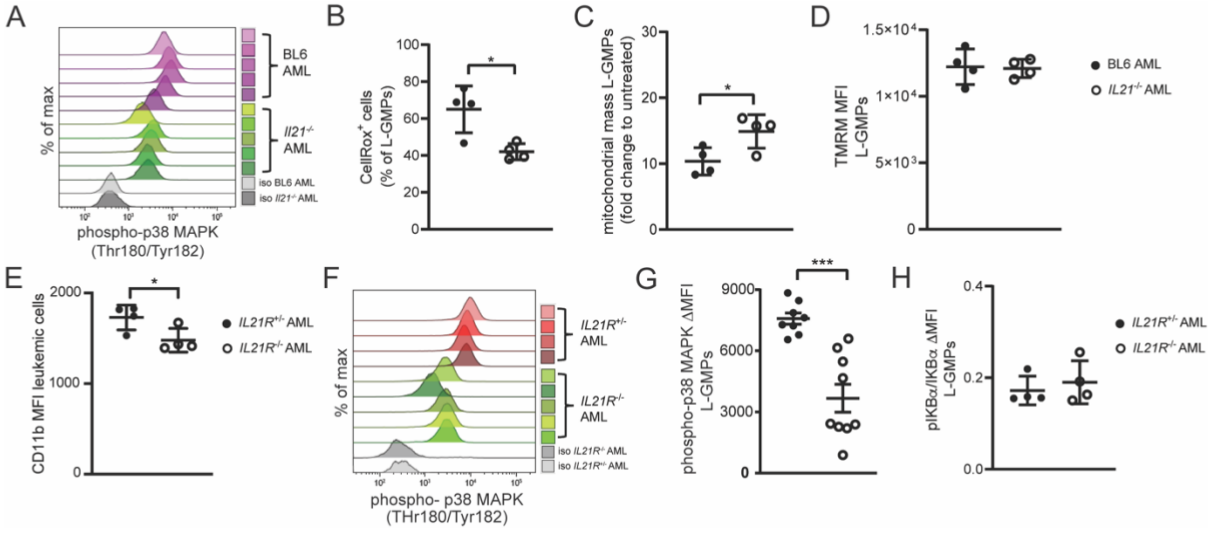


**Figure S3. Il21/Il21R signaling regulates cell L-GMPs in AML by inducing differentiation, accumulation of ROS and activation of the p38-MAPK signaling pathway.**

(**A)** Histograms showing phosphorylation of p38 MAPK (phospho-p38 MAPK) in L-GMPs from BM of BL/6 and *Il21*$^{-/-}$ AML mice (n = 5 mice/group). **(B)** Intracellular reactive oxygen species measured as frequency of L-GMPs positive to CellRox™ staining, (**C)** mitochondrial mass determined by MitoTracker™ staining and (**D)** mitochondrial membrane potential determined by TMRM™ staining of L-GMPs from BM of BL/6 and *Il21*$^{-/-}$ AML mice. One representative of two independent experiment is shown (n = 4 mice/group). Data are displayed as mean ± SD. Statistics were determined by Student's *t* test. **(E)** CD11b mean fluorescence intensity of MLL-AF9-GFP$^{+}$ leukemic cells from BM of *Il21R*$^{+/-}$ and *Il21R*$^{-/-}$ AML mice (n = 4 mice/group). Data are displayed as mean ± SD. Statistics were determined by Student's *t* test. **(F)** Histograms showing phosphorylation of p38 MAPK (phospho-p38 MAPK) and **(G)** geometric mean fluorescence intensity (MFI) quotient of phospo-p38 MAPK staining versus its isotype control on L-GMPs from BM of *Il21R*$^{+/-}$ and *Il21R*$^{-/-}$ AML mice. Two pooled independent experiments are shown (n = 4- 5 mice/group). Data are displayed as mean ± SD. Statistics were determined by Student's *t* test. **(H)** NF-kB pathway activation measured as ratio between protein expression of IκBα and its phosphorylated form pIκBα in L-GMPs from BM of *Il21R*$^{+/-}$ and *Il21R*$^{-/-}$ AML mice (n = 4 mice/group). Data are displayed as mean ± SD. Statistics were determined by Student's *t* test.

*, P < 0.05; **, P < 0.01; ***, P < 0.001.

Abbreviations: L-GMPs, leukemic granulocyte-macrophage progenitors; TMRM, tetra-methylrhodamine, methyl ester.

### Figure S4

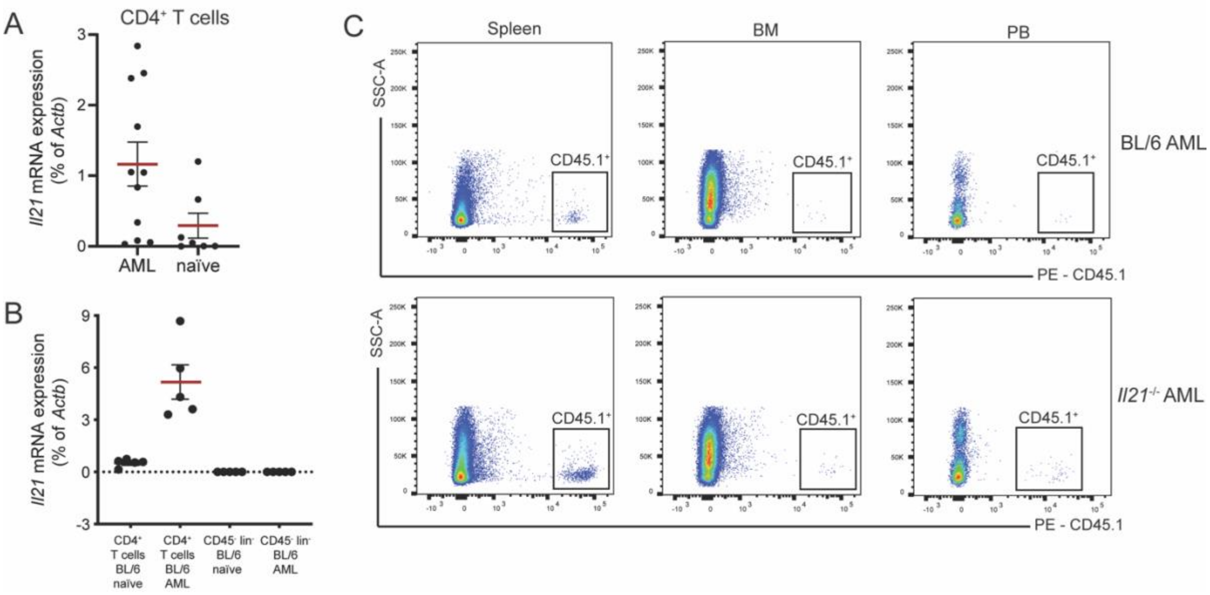


**Figure S4. CD4$^+$ T cells from AML mice express *Il21*, unlike BM stromal cells and CD4$^+$ T cells from naïve BL/6 mice. Adoptively transferred CD4$^+$ T cells can be detected by flow cytometry in spleen, BM and peripheral blood of AML mice.**

**(A)** *Il21* mRNA expression measured by qRT-PCR in FACS-sorted CD4$^+$ T cells from the BM of AML (n = 11) and naïve (n = 7) mice. Data are displayed as mean ± SEM. **(B)** *Il21* mRNA expression measured by qRT-PCR in FACS-sorted CD4$^+$ T cells and CD45$^-$lin$^-$ stromal cells from the BM of AML (n = 5) and naïve (n = 5) mice. Data are displayed as mean ± SEM. **(C)** Representative FACS plots of adoptively transferred CD45.1$^+$ cells (pre-gated on single cells) detected by flow cytometry 36 days after the transfer, in spleen, BM and PB of BL/6 and *IL21*$^{-/-}$ AML mice.

Abbreviations: PB, peripheral blood; lin, lineage.

**Figure S5**

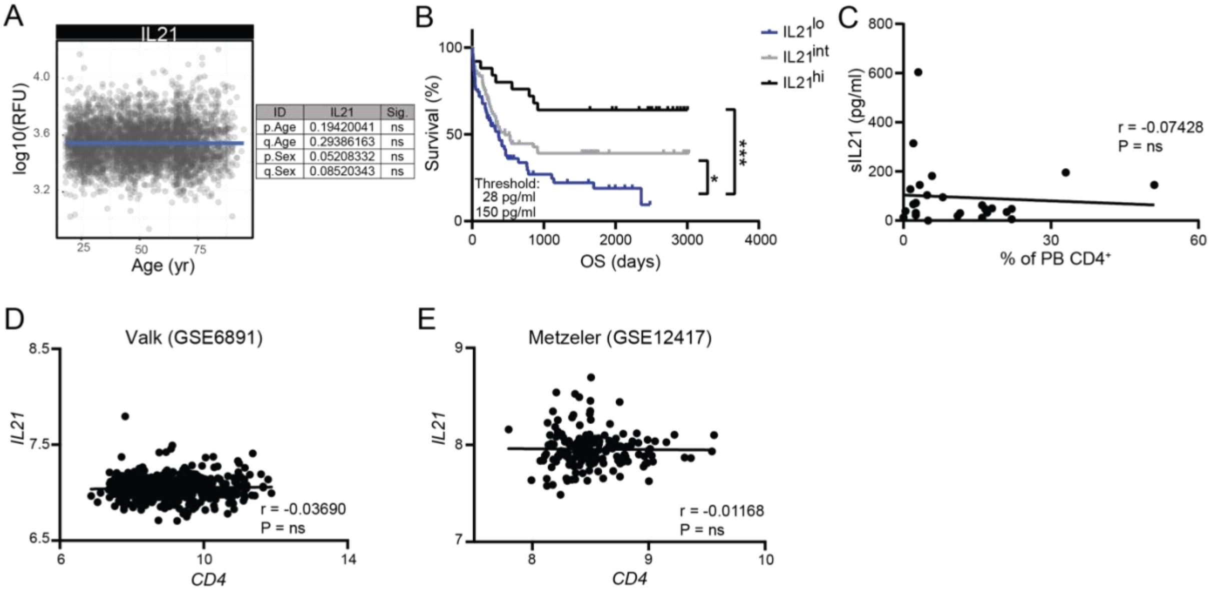


**Figure S5. sIL21 levels are not altered with age in healthy individuals and do not correlate with frequencies of $CD4^+$ T cells in peripheral blood of AML patients.**

**(A)** A publicly available plasma proteome dataset (human INTERVAL and LonGenity dataset, accession number EGAS00001002555) was analyzed for IL21 expression across lifespan (n = 4263 individuals). Statistics were determined with an age- and sex- adjusted linear model as described in Lehallier et al., 2019. **(B)** Kaplan-Meier survival curves of the entire AML patients' cohort (n = 193) divided into three groups at the sIL21 threshold of 28 pg/ml and 150 pg/ml. Statistics were determined by log-rank test. **(C)** sIL21 levels were correlated with the frequency of $CD4^+$ T cells in the peripheral blood of newly diagnosed AML patients (n = 27) determined by flow cytometry. Statistics were determined by Pearson r test. **(D, E)** *IL21* mRNA expression levels were correlated to *CD4* mRNA expression levels in the publicly available **(D)** Valk dataset (accession number GSE6891) and **(E)** Metzeler dataset (accession number GSE12417). Statistics were determined by Pearson r test.

*, P < 0.05; ***, P < 0.001.

Abbreviations: OS, overall survival; yr, years; PB, peripheral blood.

**Figure S6**

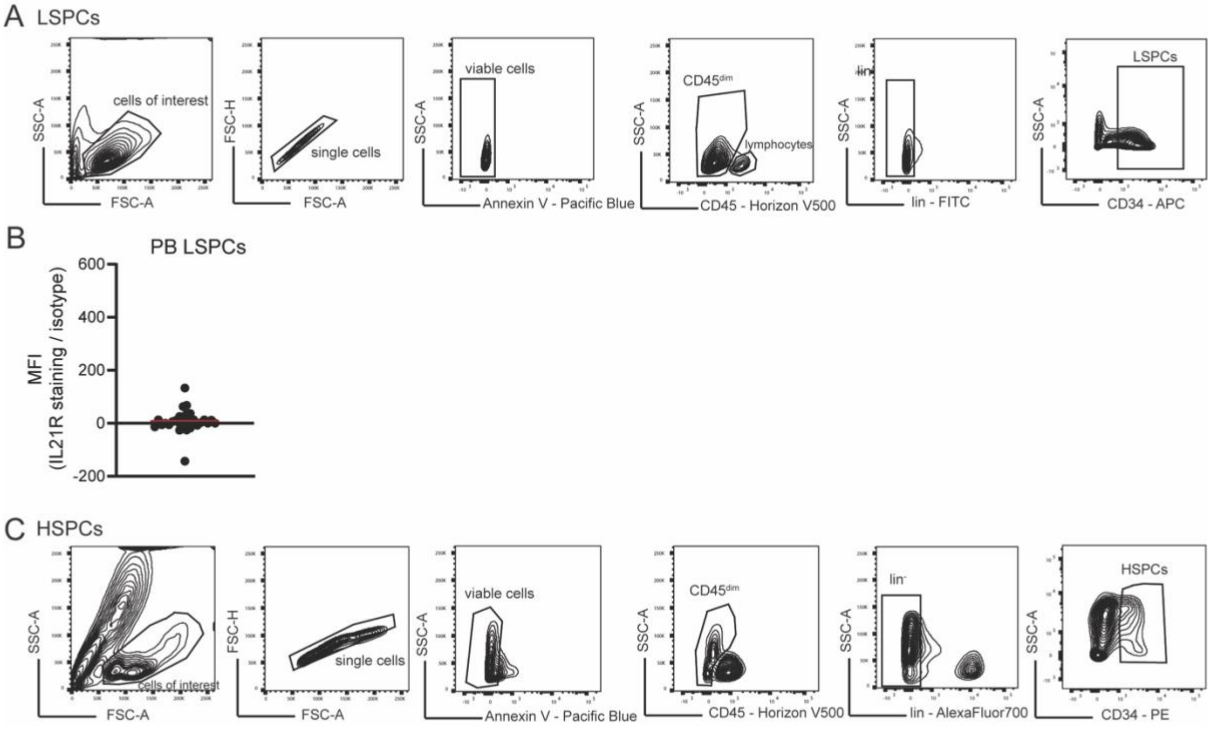


**Figure S6. FACS gating strategy and IL21R expression on LSPCs in peripheral blood.**

**(A)** Gating strategy to identify $CD45^{dim}$ $SSC^{lo}$ $lin^{-}CD34^{+}$ AML stem and progenitor cells in BM samples from newly diagnosed AML patients. **(B)** Mean fluorescence intensity (MFI) quotient of IL21R staining versus its isotype control on LSPCs (n = 30) from blood samples of newly diagnosed AML patients. Red bar indicates the mean. **(C)** Gating strategy to identify $CD45^{dim}$ $SSC^{lo}lin^{-}CD34^{+}$ stem and progenitor cells in BM samples of healthy controls who underwent BM biopsy for reason other than leukemia.

Abbreviations: LSPCs, leukemic stem and progenitor cells; PB, peripheral blood; MFI, mean fluorescence intensity; HSPCs, hematopoietic stem and progenitor cells.

**Figure S7**

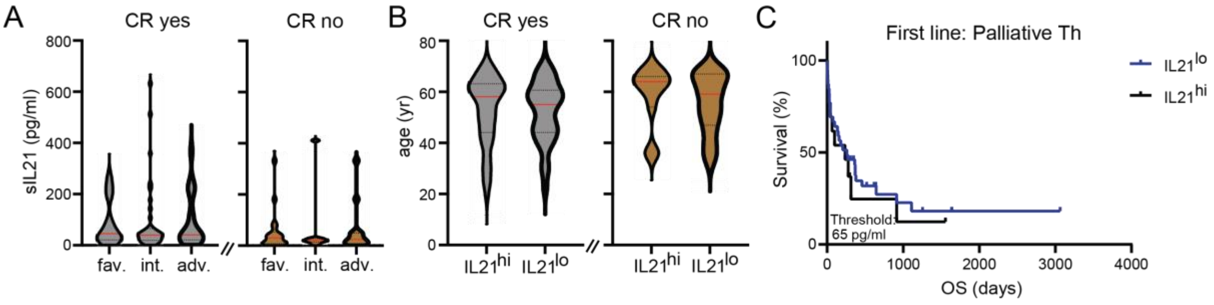


**Figure S7. sIL21 levels are not influenced by risk group and age of patients undergoing intensive chemotherapy as first line therapy. sIL21 has no prognostic value for patients undergoing palliative first line therapy.**

**(A)** sIL21 of patients that achieved and did not achieve CR, differentiated according to cytogenetic/molecular risk groups. Data are shown as mean ± SD. Statistics were determined by one-way ANOVA. **(B)** Age of patients that achieved and did not achieve CR differentiated according to sIL21 levels at the threshold of 35 pg/ml. Data are shown as mean ± SD. Statistics were determined by Mann-Whitney test. **(C)** Kaplan-Meier survival curves of the AML patients that received palliative first line therapy (n = 52) divided into two groups at the sIL21 threshold of 65 pg/ml. Statistics were determined by log-rank test.

Abbreviations: CR, complete remission; fav., favorable; int., intermediate; adv., adverse; OS, overall survival; Th, therapy.

### SUPPLEMENTARY TABLES

#### Table S1. Characteristics of AML patients.

sIL21, age at diagnosis, sex, risk category, percentage of PB blasts and BM infiltration, cytogenetic aberration and molecular diagnosis, immunophenotype and FAB classification are listed for each patient included in the study.

Abbreviations: sIL21, serum IL21; PB, peripheral blood, BM, bone marrow, infiltr., infiltration; fav., favorable; int., intermediate; adv., adverse.; n.a., not available; sec., secondary; MDS/MPS, myelodysplastic syndrome-myeloproliferative neoplasms.

| ID | sIL21 (pg/ml) | Age at diagnosis | Sex | Risk | PB blasts (%) | BM infiltr. (%) | Cytogenetics | Molecular diagnosis | Immuno-phenotype | FAB |
|---|---|---|---|---|---|---|---|---|---|---|
| 1 | 2.40 | 50 | f | adv. | 60 | 80 | Monosomy 7, Rearrangement KMT2A (MLL) | JAK 2 mutated | CD34, HLA-DR, CD13 (weak), CD33, CD64, CD117, CD38 (partial/weak). | sec. AML therapy related |
| 2 | 4.38 | 75 | f | fav. | 72 | 97 | normal | Isolated NPM1 mutation | CD13, CD33, CD38, CD4, CD64, CD16, CD71, partial CD117, cytoplamatic MPO and lysozyme. | AML-M1 |
| 3 | 4.43 | 68 | f | adv. | 42 | 40 | Complex monosomal karyotype | | | sec. AML from MDS/MPS |
| 4 | 4.90 | 70 | m | int. | 28 | 80 | Trisomy 13 | | CD34, CD38, HLA-DR, TdT, CD13; aberrant CD4. | AML-M0 |
| 5 | 6.50 | 71 | f | adv. | 26 | 40 | Deletion 5q | | n.a. | sec. AML from MDS/MPS |
| 6 | 8.10 | 39 | m | int. | 90 | 95 | t(11,17)(q23, q12,21), MLL-Rearrangement | EVI-1 | CD11b, CD16, CD38, CD56, HLA-DR, CD13, CD15, CD33, CD64, lysozyme, aberrant CD4. | AML-M4 |
| 7 | 9.30 | 59 | f | int. | 0 | 80 | normal | n.a. | n.a. | AML-M0 |
| 8 | 11.10 | 79 | f | int. | 9 | 70 | idic(X)(q13)(5)/47, sl, +idic(X) | IDH-1 mutated | CD34, CD45, HLA-DR, CD13, CD33 (partial), CD117, CD105(partial) and CD123. | AML-M2 |
| 9 | 11.78 | 40 | m | fav. | 53 | 90 | t(15,17)(q24,q21); t(13,18) unclear | t(15,17) L-Form | MPO, CD33, weak CD15 and lysozyme. | AML-M3 |

| | | | | | | | | | | |
|---|---|---|---|---|---|---|---|---|---|---|
| | | | | | | | status (possibly constitutive) | | | |
| 10 | 12.41 | 65 | f | fav. | 56 | 90 | normal | NPM1 mutated | MPO, CD33, CD15, lysozyme and partial CD68. | sec. AML therapy related |
| 11 | 13.10 | 79 | m | fav. | 23.5 | n.a. | t(15,17) | | n.a. | AML-M3 |
| 12 | 13.60 | 46 | f | fav. | 73 | 70 | t(8,21) (q22,q22), -X | | CD34, CD38, CD45, CD56, HLA-DR, CD13, CD117, MPO. | AML-M2 |
| 13 | 13.80 | 49 | f | adv. | 77 | 97 | normal | FLT3-ITD | CD34, CD38, HLA-DR, c-kit, CD13, CD33, MPO, CD16, CD64 (partial) and CD71; partially aberrant CD4. | AML-M1 |
| 14 | 13.80 | 73 | m | fav. | 20 | 50 | 47,XY,+8 | NPM1 mutated | CD38, partial HLA-DR, CD13; CD33, CD64, CD15, CD11b, CD56, MPO and lysozyme; CD14 on ca. 50% of cells. | AML-M5 |
| 15 | 15.55 | 75 | m | fav. | 0 | 60 | normal | DNMT3A and U2AF1 mutated | CD33, CD34 and c-kit. | sec. AML from MDS/MPS |
| 16 | 15.55 | 30 | f | fav. | 72.5 | 40 | normal | CEPBA mutated | CD38, HLA-DR, c-kit, CD33, CD64, CD71, MPO and CD7. CD34 strongly expressed on on ca. 65% of blasts, CD2 on ca. 65% of blasts. CD2 and CD7 aberrant expression. | AML-M1 |
| 17 | 15.55 | 56 | m | int. | 53 | 90 | normal | | CD16, CD36, CD38, CD45, CD71, CD33, CD117; aberrant CD4, CD7. | AML-M0 |
| 18 | 15.55 | 83 | f | adv. | 87.5 | 95 | Complex karyotype with Monosomy 17, unbalanced translocation Chr. 5 and 5q- deletion | | CD34, HLA-DR, CD38, CD117, CD33, CD13, CD4, CD15, CD16, CD71, cytoplasmatic TdT, MPO and lysozyme. Partially pathologic co-expression of CD56. | AML-M0 |
| 19 | 16.26 | 73 | m | n.a. | 0 | 90 | Bcl-2 and Pax-5 mutated | | CD34, CD117, TdT, CD99, CD43, MPO, aberrant BCL-2, PAX-5. | AML-M1 |
| 20 | 17.11 | 43 | f | int. | 14 | 40 | normal | RUNX-1 mutated | TdT, HLA-DR, CD33, CD34, CD38, CD45, | AML-M0 |

| | | | | | | | | | | |
|---|---|---|---|---|---|---|---|---|---|---|
| | | | | | | | | | CD13, CD117; aberrant CD4. | |
| 21 | 17.47 | 59 | m | int. | 37 | 20 | normal | n.a. | n.a. | AML-M6 |
| 22 | 17.60 | 61 | f | int. | 6 | 80 | t(15,17); MLL, EVI1; Monosomy 5+7 | PML-RARA (L-Form); FLT3-ITD | CD38, c-kit, CD13, CD33, CD56, CD71, MPO and aberrant CD2. | AML-M3 |
| 23 | 18.18 | 76 | m | int. | 18.5 | n.a. | Complex | CALR Gene Mutation Type2 | | sec. AML from MDS/MPS |
| 24 | 18.50 | 70 | m | adv. | 80 | 80 | Complex monosomal karyotype with Monosomyn 5q,7 14, 15, 16, 18 | | CD34, HLA-DR, CD117, CD13, CD33 and partial CD4. | sec. AML therapy related |
| 25 | 19.22 | 58 | m | int. | 92 | 90 | normal | FLT3-ITD | n.a. | AML-M5 |
| 26 | 19.40 | 28 | f | fav. | 74 | 90 | t(X;19)(q?22;q?12); t(15;17)(q24;q21) | PML-RARA | cytoplasmatic MPO, CD33, CD38, CD13, CD117, CD56, CD64, CD15, cytoplasmatic lysozyme and CD71. | AML-M3 |
| 27 | 19.78 | 29 | f | fav. | 0 | 95 | t(15;17)(p11.2;q21; q24) | t(15,17) S Form; FLT3-ITD | MPO, CD33 and CD15. | AML-M3 |
| 28 | 19.82 | 64 | f | adv. | 11 | 30 | Complex karytype; monosomy 5q31.2/5q33.1 | p53 Mutation | CD33, CD38, CD16, CD13, CD11b, CD71 CD64, MPO and lysozyme. | sec. AML therapy related |
| 29 | 20.84 | 79 | m | adv. | 45 | 80 | Deletion(7q) | | cytoplasmatic MPO, HLA-DR, cytoplamatic lysozyme, CD56 (bright), CD11b, CD13, CD14, CD16, CD33, CD38 and CD64. | AML-M4 |
| 30 | 21.19 | 61 | m | adv. | 4 | 10 | Complex monosomal karyotype (-7, -20, -22) | | CD34, CD45, HLA-DR, CD13, CD33, CD117, partial MPO. | sec. AML from MDS/MPS |
| 31 | 21.20 | 76 | m | adv. | 50 | 70 | normal | FLT3-ITD | n.a. | AML-M1 |
| 32 | 21.50 | 43 | m | int. | 13 | n.a. | normal | FLT3-ITD | n.a. | sec. AML from MDS/MPS |
| 33 | 21.98 | 39 | m | int. | 90 | 95 | t(11,17)(q23, q12,21), MLL-Rearrangement | EVI-1 | CD11b, CD16, CD38, CD56, HLA-DR, CD13, CD15, CD33, CD64, lysozyme, aberrant CD4. | AML-M4 |
| 34 | 22.30 | 55 | f | fav. | 99 | 95 | normal | NPM1 mutated | CD16, CD38, CD56, CD13, CD33, partial MPO. | AML-M1 |
| 35 | 22.33 | 68 | m | adv. | 33 | 20 | Monosomyn 7 (total), 5q31.2 and 5q33 | EVI 1 | CD34, c-kit. | sec. AML from MDS/MPS |
| 36 | 23.75 | 31 | m | int. | 49 | 60 | normal | FLT3-ITD | CD13, CD33, CD117, CD71, MPO, lysozyme, | AML-M4 |

| | | | | | | | | | | |
|---|---|---|---|---|---|---|---|---|---|---|
| | | | | | | | | | HLA-DR, CD16, CD64, CD38, CD4. | |
| 37 | 23.92 | 71 | m | adv. | 55 | 60 | Deletion(5)(q1?5;q3?4) | | CD34, CD38, HLA-DR, c-kit, CD33, MPO, lysozyme and CD71. | AML-M2 |
| 38 | 25.64 | 80 | f | adv. | 15.5 | 80 | normal | DNMT3A-Mutation, RUNX1-Mutation, WT1-Mutation; FLT3-ITD | CD34, HLA-DR, MPO, CD117. | sec. AML therapy related |
| 39 | 26.20 | 47 | m | int. | 27.5 | 25 | | | CD13, CD33, CD34, CD38, HLA-DR and c-kit. | sec. AML from MDS/MPS |
| 40 | 26.36 | 64 | m | adv. | 23 | 60 | Deletion(16), +8(q24), +11(q14) | | CD34, c-kit, CD13, partial HLA-DR, CD33, CD36, CD56, CD15 and weak CD105. | sec. AML from MDS/MPS |
| 41 | 27.39 | 68 | f | fav. | 2 | 70 | normal | NPM1 mutated | CD38, HLA-DR, CD13, CD71, MPO, partial CD34, c-kit, CD33, CD14, CD64, CD56 and lysozyme. | AML-M5 |
| 42 | 28.17 | 71 | m | fav. | 34 | 90 | inv(16) | | CD34, CD117, HLA-DR, cytoplasmatic MPO, CD13, CD33, CD15 and CD123. | AML-M4 |
| 43 | 28.72 | 44 | f | int. | 27 | 90 | normal | NPM1 mutated; FLT3ITD | cytoplasmatic MPO, CD13, CD117, HLA-DR and CD33. | AML-M4 |
| 44 | 29.21 | 49 | m | int. | 87 | 80 | t(8,21)(q22,q22) | | CD34, CD38, CD56, HLA-DR, CD13, CD117, MPO. | AML-M2 |
| 45 | 29.23 | 54 | f | adv. | 61 | 90 | normal | NPM1 mutated; FLT3ITD | | AML-M1 |
| 46 | 29.86 | 47 | f | adv. | 0 | 15 | Deletion(7q), inv (3q) | DVI-1 overerexpression | CD34, CD117, CD33, HLA-DR, MPO, lysozyme. | sec. AML from MDS/MPS |
| 47 | 32.12 | 46 | f | int. | 52 | 90 | Three independent clones :Trisomy 8;Deletion 7q; t(1;7)(q10;p10) | JAK 2 mutated | CD34, CD13, CD33, CD96, CD4, CD117, partial MPO and CD7. | sec. AML from MDS/MPS |
| 48 | 32.61 | 59 | f | int. | 0 | 40 | normal | FLT3 ITD | CD38, c-kit, CD15. | AML-M2 |
| 49 | 32.83 | 44 | m | adv. | 96 | 70 | Complex monosomal karyotype (-7, -9, Trisomy 8) | IDH1 mutated | Erythroid elements expressing CD71, Glycophorin A, CD36; myeloid blasts expressing CD33, CD13, | AML-M6 |

| | | | | | | | | | | |
|---|---|---|---|---|---|---|---|---|---|---|
| | | | | | | | | | CD38, HLA-DR, MPO. | |
| 50 | 33.58 | 57 | m | adv. | 17 | 30 | ERG1-Amplification (+21q22) | SRSF-2 mutated | CD34,CD45, CD15, CD117, HLA-DR, CD13, CD33, CD105, CD71, CD56, CD123, lysozyme and cytoplasmatic MPO. | sec. AML from MDS/MPS |
| 51 | 33.86 | 74 | m | int. | 20 | n.a. | n.a. | n.a. | n.a. | sec. AML from MDS/MPS |
| 52 | 34.44 | 23 | f | fav. | 15 | 60 | normal | NPM1 mutated | MPO, CD117, HLA-DR, CD13, CD123, CD15. | AML-M2 |
| 53 | 35.24 | 31 | m | int. | 49 | 60 | normal | FLT3-ITD; NPM1 mutated | CD13, CD33, CD117, CD71, MPO, lysozyme, HLA-DR, CD16, CD64, CD38, CD4. | AML-M4 |
| 54 | 35.58 | 64 | m | fav. | 50 | 70 | inv (16)(p13.1q22; +8 | IDH 2 mutated | CD45, CD34, MPO, CD117, CD13/33, HLA-DR, CD123. | AML-M4 |
| 55 | 36.74 | 76 | f | int. | 86 | 70 | Complex anomalies: deletion(5q), -6, +8, -15, -16, -17, +4mar | | cytoplasmatic MPO (partial), CD34, CD117, HLA-DR, CD13 (weak), CD33, CD71, CD15 (partial), lysozyme (weak/partial) and CD123. | AML-M2 |
| 56 | 37.17 | 49 | m | fav. | 45 | 80 | normal | IDH1 mutated | CD34, CD117, CD13/33, HLA-DR, CD35. | AML-M4 |
| 57 | 37.28 | 71 | f | adv. | 33 | 25 | Complex anomalies | p53 Mutation | MPO, CD34, CD117, HLA-DR, CD13, CD33, CD105, CD56, CD123. | AML-M2 |
| 58 | 37.40 | 69 | m | fav. | 9 | 90 | normal | NPM1 mutated | HLA-DR, CD38, CD56, CD13 (partial), CD11b, CD14, CD64, CD15, MPO, lysozyme and CD7. | AML-M5 |
| 59 | 38.19 | 28 | m | int. | 8 | 40 | normal | | CD38, CD13, CD15, CD33, CD71, HLA-DR, MPO. | AML-M2 |
| 60 | 38.55 | 71 | m | int. | 8 | n.a. | normal | JAK 2 mutated | CD34, HLA-DR, CD117, CD13, CD33 and MPO. | sec. AML from MDS/MPS |

| | | | | | | | | | | |
|---|---|---|---|---|---|---|---|---|---|---|
| 61 | 39.63 | 63 | f | adv. | 99 | 90 | normal | NPM1 mutation type A , FLT3-ITD | MPO, CD15, CD33, HLA-DR, c-kit, CD34. | AML-M1 |
| 62 | 39.92 | 72 | m | adv. | 20 | 80 | inv(9)(p11q13) | NPM1, IDH-2 and SRSF-2 mutated | CD56, CD33, CD14, CD163. | sec. AML from CMML, AML-M4 |
| 63 | 40.74 | 60 | m | adv. | 10 | 35 | Complex anomalies: -3, -5, -7, +8, -12, deletion(13), -17, -18, +3mar | TP53 Mutation | CD34, CD117, CD13, CD33, HLA-DR, MPO, CD123. | AML-M4 |
| 64 | 41.26 | 57 | m | int. | 57 | 95 | t(9,11)(p21;q23) | U2AF1 mutated | HLA-DR, CD117, CD33, CD68, CD15, CD4 and lysozyme. | AML-M5 |
| 65 | 42.90 | 86 | m | int. | 64 | n.a. | n.a. | n.a. | HLA-DR, CD38, CD34, CD117, cytoplasmatic MPO, CD33, CD16, CD71, lysozyme and aberrant CD7. | AML-M1 |
| 66 | 43.90 | 40 | f | int. | 20 | 80 | t(8,21), Deletion 9q | ASXL1 mutated | cytoplasmatic MPO, HLA-DR, CD34, CD38, CD13, CD15, CD117 and CD56. | AML-M2 |
| 67 | 44.07 | 78 | f | adv. | 6 | 40 | Complex anomalies | | CD34, c-kit, CD33. | AML-M2 |
| 68 | 44.20 | 81 | m | adv. | 96 | 90 | Deletion -9, der(11)t(9.11)(q13, q?22); monosomy 11q23.3 | | CD13, CD33, CD15, MPO, CD7, CD2 (partial), CD64 (partial, c-kit, CD34 (partial) and HLA-DR. | AML-M1 |
| 69 | 44.89 | 63 | f | int. | 7 | 50 | normal | IDH2 Mutation | CD34,CD117, CD33, CD16, CD71, cytoplasmatic MPO; partial HLA-DR and CD38; partial CD7 co-expression. | AML-M2 |
| 70 | 45.07 | 50 | m | fav. | 85.5 | 70 | normal | CEPBA mutated | CD34, HLA-DR, CD117, aberrant CD7, CD15 and MPO; weak CD13 and CD33. | AML-M1 |
| 71 | 45.26 | 69 | f | adv. | 11 | 70 | normal | FLT3-ITD; TET2 and DNMT3A mutated | CD13, CD33, HLA-DR, CD123 and MPO. | AML-M5 |
| 72 | 45.49 | 69 | m | adv. | 99 | 80 | Trisomy 11q23.3 | | CD117, MPO, CD13, CD33, CD34, CD16, CD11b, CD38, CD71, CD4, HLA-DR. | AML-M2 |
| 73 | 46.34 | 65 | m | adv. | 94 | 85 | t(8,21) | FLT3-ITD | cytoplasmatic MPO (weak/partial), cTdT, CD34, CD38, CD13, | AML-M2 |

| | | | | | | | | | | |
|---|---|---|---|---|---|---|---|---|---|---|
| | | | | | | | | | CD33, CD117, CD123, CD11b, HLA-DR, CD19, CD4 and CD2 (weak/partial). | |
| 74 | 46.44 | 51 | m | int. | 90 | 90 | normal | FLT3-ITD, NPM1 mutated | c-kit, MPO, CD33, HLA-DR (partial), CD68, lysozyme (partial), CD34, CD117, CD13, HLA-DR, MPO, CD7 (aberant), CD15, CD123. | AML-M1 |
| 75 | 48.00 | 20 | f | int. | 38 | 60 | normal | n.a. | CD16, CD34, CD38, CD45, CD71, HLA-DR, CD13, CD33, CD117, MPO. | AML-M2 |
| 76 | 49.83 | 25 | f | fav. | 32.5 | 60 | t(8,21), -X | | CD13, CD117, MPO, HLA-DR, CD34. | sec. AML therapy related, M1-2 |
| 77 | 50.90 | 63 | m | int. | 23 | 50 | inv(7)(q22q36) | RUNX1 mutated | CD45, CD13, CD33, CD117, CD34, MPO, CD7. | AML-M4 |
| 78 | 51.27 | 80 | f | adv. | 40 | 25 | normal | RUNX-1 mutated | | sec. AML from MDS/MPS |
| 79 | 52.60 | 69 | f | int. | 1 | 90 | normal | FLT3-ITD; WT1 mutated | HLA-DR, CD117, CD13, partial CD34 and CD71. | AML-M5 |
| 80 | 56.20 | 64 | m | fav. | 2 | 30 | Deletion(5)(q21-q33) | RUNX1 mutated | CD34, CD38, CD45, CD56, HLA-DR, CD13, CD33, CD117. | sec. AML from MDS/MPS |
| 81 | 56.57 | 66 | f | adv. | 36 | n.a. | Complex anomalies: -3,Deletion(5)(q13q33),+8,-11,Deletion(12)(p12p13),-18 | | CD34, CD38, CD117, HLA-DR, CD13, CD33, MPO, lysozyme, CD71 and weak aberrant CD7. | sec. AML therapy related, M0 |
| 82 | 59.60 | 69 | m | int. | 90 | 80 | normal | n.a. | HLA-DR, CD33. | AML-M4 |
| 83 | 61.87 | 51 | f | adv. | 54 | 70 | normal | EVI1 overexpression, RUNX1 mutated, FLT3-ITD | CD33, CD68, c-kit and CD15; partial CD34, CD36,CD45, HLA-DL, CD13, MPO. | AML-M4 |
| 84 | 62.13 | 65 | f | fav. | 54 | n.a. | normal | IDH2 mutated | CD13, CD15, CD33, CD64, CD11b, CD36, CD38, CD56 and HLA-DR; cytoplasmatic MPO and | AML-M4 |

| | | | | | | | | | | |
|---|---|---|---|---|---|---|---|---|---|---|
| | | | | | | | | | lysozyme; partial CD16 and CD4. | |
| 85 | 62.51 | 45 | m | adv. | 96 | 90 | normal | IDH1mutated; FLT3-ITD | | AML-M1 |
| 86 | 63.67 | 44 | f | int. | 27 | 90 | normal | NPM1, FLT3 mutated | cytoplasmatic MPO, CD13, CD117, HLA-DR and CD33. | AML-M4 |
| 87 | 65.93 | 77 | m | int. | 0 | 50 | normal | ASXL1, IDH2 mutated | MPO, CD34, CD117, CD13, HLA-DR, CD33. | AML-M2 |
| 88 | 68.37 | 62 | f | int. | 7 | 50 | normal | IDH2 mutated | CD34,CD117, CD33, CD16, CD71, cytoplasmatic MPO; partial HLA-DR and CD38; partial CD7 co-expression. | AML-M2 |
| 89 | 72.58 | 34 | m | fav. | 82 | 95 | inv (16)(p13.1q22) | c-kit-mutated | cytoplasmatic MPO, HLA-DR, CD34, CD13, CD33, CD117, CD123, CD15 (weak, partial). | AML-M4 |
| 90 | 74.80 | 69 | f | n.a. | 25 | n.a. | n.a. | n.a. | CD11b, CD14, CD64, CD13, CD33, CD4, CD34, CD36, CD56, HLA-DR, MPO and lysozyme. | AML-M4 |
| 91 | 78.77 | 67 | f | adv. | 51 | 25 | inv(3)(q21q26), -7, Deletion(11) | EVI1 mutated | CD34, CD13, CD14, CD33, HLA-DR. | sec. AML from MDS/MPS |
| 92 | 80.82 | 58 | m | fav. | 46 | 85 | normal | NPM1 mutated | CD117, CD13, CD33. | AML-M4 |
| 93 | 81.81 | 63 | m | fav. | 58 | 90 | normal | NPM1, IDH1 mutated | CD117 (partial), CD33, CD123, MPO, lysozyme. | AML-M1 |
| 94 | 81.95 | 73 | f | fav. | 90 | 80 | normal | NPM1 mutated | CD14, CD68, CD11c, CD33, HLA-DR. | AML-M5 |
| 95 | 82.28 | 70 | m | int. | 47 | 90 | normal | CEPBA, IDH-2 mutated | MPO, CD33, CD34, HLA-DR and c-kit. | AML-M1 |
| 96 | 89.25 | 71 | m | adv. | 90 | 20 | normal | ASXL1 and IDH2 mutated | CD34, CD117, CD13, HLA-DR and MPO; weak/partial expression of CD33, CD123 and lysozyme. | AML-M2 |
| 97 | 89.90 | 62 | f | fav. | 83 | 80 | Deletion(7)(q22) | NPM1, IDH1 mutated | CD13, CD33, CD15, MPO, CD64 (partial), lysozyme, CD71, c-kit and partial CD34. | AML-M2 |

| | | | | | | | | | | |
|---|---|---|---|---|---|---|---|---|---|---|
| 98 | 90.25 | 74 | f | int. | 20 | 30 | normal | NPM1 mutated, FLT3-ITD | CD117, CD13, CD33, CD38, CD11b, CD15, HLA-DR and partial CD14. | AML-M4 |
| 99 | 106.18 | 48 | f | adv. | 10 | 90 | normal | RUNX1 mutated | cytoplasmatic MPO (partial), CD34, CD117, CD13, HLA-DR, CD71, cTdT, CD15 (partial), aberrant CD4, CD12. | AML-M1 |
| 100 | 118.79 | 70 | f | adv. | 57 | 60 | Complex anomalies, with -7 and -5 | | CD13, CD33, MPO, CD123, CD34, c-kit, HLA-DR, CD105 and CD11b. | AML-M2 |
| 101 | 119.31 | 25 | f | adv. | 10 | 25 | Rearrangement KMT2A(MLL); t(11,19)(q23,q13.1) | EVI-1 overexpress ion | CD34 (partial), CD38, HLA-DR, c-kit and CD33. | AML-M4 |
| 102 | 141.50 | 46 | f | int. | 52 | 90 | Three independent clones:Trisomy 8; deletion 7q; t(1;7)(q10;p10) | JAK2 mutated | CD34, CD13, CD33, CD96, CD4, CD117, MPO and CD7. | sec. AML from MDS/MPS |
| 103 | 176.24 | 52 | m | int. | 8 | 15 | normal | FLT3-ITD | CD34, CD38, CD117, HLA-DR, CD13, CD33, MPO and aberrant CD4. | AML-M6 |
| 104 | 179.08 | 69 | f | fav. | 25 | 40 | normal | NPM1 mutated | CD45, HLA-DR, CD117 (partial), CD13/33, CD14, CD64, IREM-2 and CD35. | AML-M4 |
| 105 | 192.34 | 58 | f | fav. | 82 | 90 | normal | NPM1, IDH1 R132 mutated | CD33, MPO, c-kit and CD123. | AML-M1 |
| 106 | 199.80 | 40 | f | fav. | 47 | 85 | deletion-X, t(8,21) | | CD34, CD38, HLA-DR, c-kit, CD13, MPO, CD71 and aberrant CD19; partial CD33, CD15 and cytoplasmatic TDT. | sec. AML therapy related |
| 107 | 208.67 | 65 | f | adv. | 93 | 90 | normal | NPM1 mutated, FLT3-ITD | CD13,CD33, CD117, MPO, HLA-DR, CD34, CD38, CD71, CD11b. | AML-M1 |
| 108 | 212.90 | 66 | m | adv. | 0 | 90 | normal | FLT3-ITD | CD13, CD117, CD34, CD38, CD4 (aberrant), HLA-DR and TdT. | AML-M0 |

| | | | | | | | | | | |
|---|---|---|---|---|---|---|---|---|---|---|
| 109 | 222.32 | 44 | f | int. | 35 | n.a. | normal | NPM1 mutated, FLT3-ITD | | AML-M5 |
| 110 | 235.52 | 63 | f | adv. | 26 | 90 | normal | FLT3-ITD | CD11b, CD38, CD45, CD71, HLA-DR, CD13, CD14, CD15, CD33, CD64, lysozyme, aberrant CD4. | AML-M4 |
| 111 | 236.00 | 59 | m | adv. | 70 | 80 | Monosomy 7q31.2 | IDH2 mutated | CD34, CD33, c-kit, TdT (partial), HLA-DR, MPO and CD15. | AML-M2 |
| 112 | 239.40 | 43 | f | int. | 14 | 40 | normal | RUNX-1 mutated | TdT, HLA-DR, CD33, CD34, CD38, CD45, CD13, CD117, aberrant CD4. | AML-M0 |
| 113 | 245.98 | 68 | m | fav. | 80 | 90 | inv (16) | | CD33, CD34, CD38, CD117, HLA-DR, and partial CD13, CD15, CD64, CD4 and TdT. | sec. AML from MDS/MPS |
| 114 | 246.42 | 60 | m | fav. | 2 | 90 | normal | NPM1 mutated | CD15, HLA-DR, CD33, CD68, MPO, CD117, CD56. | AML-M4 |
| 115 | 258.74 | 80 | f | fav. | 15 | 90 | normal | NPM1 mutated | CD11b, CD13, HLA-DR. | AML-M5 |
| 116 | 267.87 | 32 | m | fav. | 95 | 90 | Complex Translocation with possibly criptic inversion 16 | NPM1 mutated | CD34, CD38, HLA-DR, CD117, CD13, CD33, MPO, CD71, partially aberrant CD4. | AML-M2 |
| 117 | 273.13 | 64 | m | fav. | 27 | 80 | t(15,17) | PML-RARA, L-Form | CD13, CD33, MPO, CD38, CD45 and CD71. | AML-M3 |
| 118 | 274.77 | 63 | f | fav. | 53 | 90 | normal | NPM1 mutated | CD16, CD38, CD45, CD71, CD13, CD33, CD117, HLA-DR, MPO. | AML-M2 |
| 119 | 332.26 | 36 | m | adv. | 29 | 25 | Complex anomalies: der(7)t(3;7)(q21;p22), der(17;18)(q10;10), der(19;20)(q10;p10, +mar(cp6)/43-44.sl,+3, der (3;5)(q10;q10), -der(7), +7(cp3) | p53 mutated | CD45, CD34, CD117, HLA-DR, CD13, CD33, CD105 and CD123. | AML-M0 |
| 120 | 347.19 | 67 | f | int. | 3 | n.a. | normal | FLT3 ITD | CD38, c-kit, CD15. | AML-M7 |
| 121 | 358.81 | 59 | f | int. | 0 | 40 | n.a. | n.a. | n.a. | AML-M2 |
| 122 | 360.30 | 65 | f | fav. | 54 | n.a. | normal | IDH2 mutated | CD13, CD15, CD33, CD64, CD11b, CD36, CD38, CD56 and HLA-DR.; cytoplasmatic MPO and | AML-M4 |

| | | | | | | | | | | |
|---|---|---|---|---|---|---|---|---|---|---|
| | | | | | | | | | lysozyme; partial CD16 and CD4. | |
| 123 | 366.87 | 40 | f | adv. | 2 | 30 | normal | EVI-1 mutated | CD13, CD33, CD117, MPO, CD34, CD38, CD71, HLA-DR, CD4 (aberrant), CD15. | AML-M1 |
| 124 | 376.92 | 45 | m | adv. | 96 | 90 | normal | IDH1 mutated, FLT3-ITD | | AML-M1 |
| 125 | 512.60 | 60 | m | int. | 16 | 80 | Two independent clones: trisomy 13, trisomy 21 | | CD34, CD38, HLA-DR, TdT, CD13, aberrant CD4. | AML-M0 |
| 126 | 632.75 | 60 | m | int. | 51.5 | 70 | normal | n.a. | CD34, CD38, HLA-DR, CD117, MPO, CD13, CD33 (partial), TdT, CD71 and lysozyme. | AML-M2 |
| 127 | 21.05 | 65 | f | fav. | 3 | 70 | normal | DNMT3A mutated | CD11b, CD45, CD71, HLA-DR, CD13, CD33, CD117, MPO, CD123. | AML-M2 |
| 128 | 17.39 | 54 | f | adv. | 85 | 90 | Deletion 7; EVI1-rearrangement | NRAS mutated | CD34, CD38, c-kit, HLA-DR, CD13, CD33, CD4, CD7 (partial), and CD56 (partial). | sec. AML therapy related |
| 129 | 5.32 | 59 | f | fav. | 68 | 85 | normal | CEPBA mutated | CD34, CD117, CD13, CD33, HLA-DR, MPO, CD7, CD56, CD71, CD15. | AML-M1 |
| 130 | 114.39 | 67 | f | adv. | 7.2 | 20 | normal | TP53 mutated, KRAS mutated | | sec. AML from MDS/MPS |
| 131 | 4.88 | 33 | m | int. | 15.5 | 25 | normal | NPM1 mutated; FLT3 mutated; NRAS mutated; DNMT3A mutated | CD117, CD13, CD33, HLA-DR, MPO, CD123, CD15 (partial), aberrant HLA-DR, CD36. | AML-M2 |
| 132 | 1.98 | 64 | m | int. | 66 | 90 | -Y | EVI overrexpression, RUNX1 mutated | CD45, CD34, CD117, CD13/33, HLA-DR, CD123. | AML-M4 |
| 133 | 9.35 | 64 | m | adv. | 0 | 80 | Trisomy 13, Trisomy 19 | RUNX1, SRSF2, PHF6 mutated | CD13, CD15, CD33, CD117, MPO, CD34, CD45, HLA-DR TdT, CD123, CD105. | AML-M0 |
| 134 | 53.36 | 66 | m | adv. | 2 | 40 | Deletion(5q) | ASXL1, IDH2 mutated | CD13, CD33, CD117, MPO, CD123, CD105, CD34. | AML-M2 |

| | | | | | | | | | | |
|---|---|---|---|---|---|---|---|---|---|---|
| 135 | 11.10 | 75 | m | adv. | 25 | 40 | Complex monosomal karyotype | TP53 mutated | n.a. | AML-M0 |
| 136 | 16.90 | 71 | m | fav. | 65 | 65 | Deletion(9)(q13q22) | NPM1 mutated, PTPN11 genic variant | Myeloid population: CD34, CD117, CD13/CD33, HLA- DR. Monocytic population: CD45, HLA-DR, CD64, IREM-2. | AML |
| 137 | 11.33 | 75 | m | adv. | 64 | 80 | 47, +13 | ASXL1 , IDH2, RUNX1, SRSF2, STAG2 mutated | CD34, CD117, CD13, HLA. | AML-M0 |
| 138 | 12.79 | 60 | m | adv. | 37.5 | n.a. | Complex karyotype | TP53 mutated | CD34, CD117, CD13, CD33, HLA-DR, CD105, CD123. | AML-M0 |
| 139 | 22.09 | 57 | m | fav. | 70 | 90 | inv(16)(p13.1 q22.1) | Two mutations in NF1 gene; rearrangem ent CBFB-MYH11 | CD34, CD117, CD13/33, HLA-DR. | AML-M4 |
| 140 | 29.44 | 70 | m | adv. | 5 | 6 | Deletion 9p and 14q | JAK2 , RUNX1, MPL, SRSF2, NRAS, TET2 mutated | CD34, CD117, CD13, CD33, HLA-DR, cytoplasmatic MPO, CD105, CD123, CD7, CD56. Monocytic population: CD14, CD64, IREM-2, CD35, CD11b, HLA-DR, aberrant CD105 and CD56. | sec. AML from MDS/MPS |
| 141 | 5.74 | 60 | f | adv. | 11 | 75 | normal | RUNX1, BCOR mutated | CD34, CD117, CD13, CD33, HLA-DR, CD7, CD105, CD123, TdT. | AML-M1 |
| 142 | 31.56 | 53 | m | int. | 81 | 75 | normal | IDH1; RUNX1; NPM1; FLT3 mutated | Myeloid population: CD117, CD13, CD33, HLA-DR, CD123. Monocytic population: CD14, CD16, IREM-2, CD35, CD36, CD64, HLA-DR, CD11b. | AML-M1 |
| 143 | 18.73 | 74 | f | adv. | 60 | 90 | normal | NPM1, TET2, ZRSR2 mutated | | AML-M4 |
| 144 | 38.36 | 54 | m | int. | 68 | 80 | normal | IDH1, IDH2, NPM1, | CD117, CD13, CD33, | AML-M5 |

| | | | | | | | | | | |
|---|---|---|---|---|---|---|---|---|---|---|
| | | | | | | | | DNMT3A mutated; FLT3-ITD | HLA-DR, CD123, CD7. | |
| 145 | 11.79 | 39 | m | fav. | 47 | 80 | t(8;21) (q21.3; q22.1) | kit mutated | CD34, CD117, CD13/CD33. | AML-M2 |
| 146 | 18.93 | 59 | m | int. | 22.1 | 95 | t(11;17)(q23;q2124) ; add(X)(q22),add(6)( q10),- 12,add(21)(q22),+m ar[6] | NRAS mutated | n.a. | AML-M1 |
| 147 | 0.03 | 70 | f | adv. | 80 | 30 | normal | ASXL1, TET2, KRAS mutated | HLA-DR, CD33, CD38, CD64, CD14, CD93, IREM-2, lysozyme. | AML-M4 |
| 148 | 16.18 | 73 | f | adv. | 41.5 | 60 | inv(16) | FLT3, ASXL1, EZH2, RUNX1 mutated; CBL, PRPF8, SETBP1 genic variant | | sec. AML from MDS/MPS |
| 149 | 150.63 | 81 | m | int. | 20.5 | 80 | normal | FLT3ITD; NPM1, FLT3, NRAS, TET2 mutated | HLA-DR, CD33,CD123, CD11b, CD35, CD64, CD14, IREM2, CD36. | AML-M4 |
| 150 | 34.49 | 69 | f | fav. | 1 | 5 | t(15;17) | PML-RARA, JAK2 mutated | CD34, CD117, CD13/CD33. | AML-M3 |
| 151 | 38.13 | 64 | m | adv. | 68 | 45 | del-7; t(9;22)(q34;q11.2) | RUNX1 mutated; TET2, PTPN11, PRPF8, NF1 gene variant; BCR-ABL1 | CD34, CD117, partial CD13, CD33, HLA-DR. | AML-M2 |
| 152 | 13.08 | 67 | m | adv. | 51.5 | 90 | inv(3)(p21-22q26),-7 | IDH2, TET2, NRAS, DNMT3A, SRSF2 mutated | CD45, CD34, CD13, CD33, CD117, HLA-DR, CD7, CD2, CD11b, CD105, CD123, CD38. | sec. AML therapy related |
| 153 | 50.52 | 68 | f | int. | 79.5 | 90 | normal | ASXL1, TET2 2 mutated | CD117, HLA-DR, CD13, CD33,CD34, CD45. | AML-M2 |
| 154 | 29.15 | 50 | f | fav. | 1.5 | 50 | add(8)(q22),t(10;15; 17)(q23;q24;q21) | | | AML-M3 |
| 155 | 4.21 | 54 | m | fav. | 3 | 25 | +22 | STAG2, NPM1 mutated | CD45, CD34, CD117, CD13, CD33, CD38, HLA-DR, MPO, CD105, CD71, CD123. | AML-M2 |
| 156 | 2.17 | 68 | f | adv. | 84 | 45 | del,-4,-5,-7,-17,-22, | TP53 mutated | CD34, CD117, HLA-DR, CD13/33. | AML-M1 |
| 157 | 12.52 | 30 | m | int. | 66.5 | 60 | t(10;11)(p12;q14) | EZH2 mutated; rearrangem ents | CD117, HLA-DR, CD34, CD33. | AML-M1 |

| | | | | | | | | | | |
|---|---|---|---|---|---|---|---|---|---|---|
| | | | | | | | | PICALM-MLLT10 | | |
| 158 | 14.49 | 75 | f | adv. | 11 | 25 | Trisomy 8, 47,XX,+8 | ASXL1, KRAS, U2AF1, SETBP1, STAG2 mutated | CD34, CD117, CD13, CD33, HLA-DR, CD105. | sec. AML from MDS/MPS |
| 159 | 0.84 | 66 | m | int. | 5 | 20 | normal | NRAS, DNMT3A, U2AF1, ETV6 mutated | CD34, HLA-DR, CD117 and CD13/33. | AML-M4 |
| 160 | 0.00 | 62 | m | int. | 19 | 90 | normal | NPM1, DNMT3A, TET2, PTPN11 mutated | CD33, HLA-DR, CD11b, CD105, CD36, CD14, CD64, CD16, CD56, IREM2. Aberrant partial CD2 and CD4. | AML-M5 |
| 161 | 9.05 | 68 | f | adv. | 81.5 | 85 | der(2)t(1;2)(q25-31;q31), der(7)t(7;8)(q31;q21) | FLT3ITD; IDH1, FLT3 mutated; DNMT3A, IKZF1, PTPN11, BCOR gene variant | CD45, CD117, CD13, CD33, HLA-DR, CD7, CD105, CD123. | AML-M0 |
| 162 | 35.38 | 62 | m | fav. | 61 | 75 | normal | IDH2 mutated, DNMT3A gene variant | MPO, CD34, CD45, CD117, HLA-DR, CD13, CD105, CD33, CD123. | AML-M4 |
| 163 | 17.58 | 64 | m | int. | 3.5 | n.a. | Deletion(4)(q31q32),+8; Trisomy 8 | kit, TET2 2 mutated | CD45, CD117 CD13, CD33, CD11b, CD36, CD56, HLA-DR. | AML-M6 |
| 164 | 12.23 | 36 | f | int. | 94 | 90 | normal | FLT3ITD; IDH2, NPM1, FLT3 mutated | CD34 CD117, CD13, CD33, HLA-DR, MPO. | AML-M1 |
| 165 | 0.00 | 38 | m | adv. | 4 | 43 | inv(3)(q21q26) | SF3B1, ETV6 mutated | Partial CD34 | sec. AML from MDS/MPS |
| 166 | 18.95 | 41 | m | adv. | 6 | 25 | normal | RUNX1 mutated; FLT3ITD | CD34, CD15, CD33, MPO, lysozyme, HLA-DR, CD2; isolated CD3, CD5, CD4, CD8. | AML unspecified |
| 167 | 105.71 | 54 | m | int. | 68 | 80 | normal | IDH1, IDH2, NPM1, DNMT3A mutated; FLT3-ITD | CD117, CD13, CD33, HLA-DR, CD123, CD7. | AML-M5 |
| 168 | 5.93 | 52 | m | fav. | 26 | 80 | normal | NPM1 mutated | CD117, HLA-DR, CD13, CD33, MPO, CD123, lysozyme, CD16 (partial), CD15 (partial). | AML-M4 |

| | | | | | | | | | | |
|---|---|---|---|---|---|---|---|---|---|---|
| 169 | 23.12 | 73 | f | adv. | 45 | 80 | Deletion.5q | IDH2, RUNX1, DNMT3A, SRSF2 mutated | MPO, CD34, CD117, CD13, CD33, HLA-DR, CD123, CD105, CD15, cTdT. | sec. AML therapy related |
| 170 | 0.00 | 79 | m | adv. | 53 | 80 | Monosomy 7 | ASXL1, TET2, KRAS, SH2B3, U2AF1 mutated | CD34 (partial), CD13, CD33, MPO, CD35, CD11b, CD14, CD15 and (partial) CD16, CD64dim, (partial) IREM-2, (partial) CD105, CD56, weak CD123, lysozyme. | sec. AML from MDS/MPS |
| 171 | 16.37 | 69 | m | int. | 62 | 90 | normal | FLT3-ITD; U2AF1, BCOR, TET2 mutated | CD117, CD13, CD33, HLA-DR, MPO, CD7, CD123, CD36 (partial). | AML-M1 |
| 172 | 203.89 | 67 | m | adv. | 8 | 10 | +8 | RUNX1 and ASXL1 mutated | CD34, CD117, CD13, CD33, HLA-DR, MPO, CD56. | AML-M4 |
| 173 | 411.10 | 54 | f | int. | 46 | 95 | normal | TET2, NPM1 mutated | MPO, CD33, CD117, CD123. | AML-M4 |
| 174 | 94.81 | 80 | m | adv. | 24 | n.a. | normal | ASXL1 , TET2, ZRSR2 mutated | CD34, CD117, CD13, CD33, HLA-DR, MPO, CD105, and aberrant CD2, CD7 (partial) and CD123. | AML-M4 |
| 175 | 66.01 | 88 | f | adv. | 16 | n.a. | + idic(X)(q13) | ASXL1, TET2, SRSF2 mutated | MPO, CD34, CD117, CD13, CD33, HLA-DR, CD11, CD14, IREM, CD4, CD123. | AML-M4 |
| 176 | 314.32 | 68 | f | adv. | 83 | 100 | normal | FLT3-ITD; TET2 mutated | MPO, CD117, HLA-DR, CD13, CD33, CD11b, CD105, CD123, CD15, CD71. | AML-M1 |
| 177 | 181.54 | 60 | m | adv. | 8 | 30 | Complex anomalies with KMT2A (MLL) amplification | TP53 mutated | CD34, CD117, HLA-DR, CD13, CD33. | AML-M2 |
| 178 | 47.67 | 57 | f | adv. | 13 | 95 | Complex anomalies: t(X;3), deletion11, Trisomyn 1, 6, 9, 11 and 12 | WT1 and BCOR mutated | CD45, CD34, CD117, HLA-DR, CD13, CD33, MPO, TdT; aberrant CD56. | AML-M2 |
| 179 | 30.17 | 71 | f | fav. | 86 | 95 | normal | FLT3-ITD; DNMT3A, NPM1 mutated | CD13, HLA-DR, MPO, CD16, CD15, CD14, IREM- | AML-M5 |

| | | | | | | | | | | |
|---|---|---|---|---|---|---|---|---|---|---|
| | | | | | | | | | 2, CD64, CD36, CD56 (partial), CD123, lysozyme. | |
| 180 | 32.60 | 49 | m | int. | 33 | 60 | der(16)t(11;16), Trisomy 14 | IDH2, DNMT3A mutated | CD45, CD34, CD117, CD13, MPO, CD105, CD123. | AML-M2 |
| 181 | 121.09 | 67 | m | int. | 58.5 | 95 | normal | IDH-2, TET2, SRSF2 mutated | CD117, CD33, MPO, CD105, CD123. | AML-M1 |
| 182 | 174.09 | 61 | m | adv. | 44 | 70 | Monosomy 7 | RUNX1; NRAS; DNMT3A; TET2; SRSF2 mutated | CD34, CD117, CD13, CD105, CD123 and MPO. | AML-M4 |
| 183 | 413.28 | 70 | m | adv. | 80.5 | 90 | Tetrasomy 13 | RUNX 1, DNMT3A, SRSF2 mutated | CD34, CD11b, CD45, HLA-DR and CD38, CD117 (partial). | AML-M5 |
| 184 | 49.35 | 70 | m | int. | 15 | 80 | +8 | FLT3 ITD; IDH2, kit mutated, KRAS, BCOR, DNMT3A gene variant | | AML-M1 |
| 185 | 168.06 | 39 | m | adv. | 85 | 80 | +8, deletion(10)(p?11.2 p?15), add(17)(q24) | FLT3 mutated | MPO, CD34 (partial), CD117, HLA-DR, CD13 (partial), CD33, CD105, CD36 (partial). CD71, CD56, CD4, CD15, CD123. | AML-M4 |
| 186 | 195.44 | 60 | f | adv. | 39 | 95 | Complex | DNMT3A, TET2 mutated; FLT3ITD | | AML-M1 |
| 187 | 72.03 | 36 | f | fav. | 6.48 | n.a. | normal | NRAS, DNMT3A, KRAS, mutated CBFB-MYH11 | CD34, CD117, CD13/CD33. | AML, myeloid sarcoma (MS), chloroma |
| 188 | 144.70 | 74 | m | int. | 88.5 | 90 | t(3;16)(q?21;q?22), +8, t(2;7)(p21;q36)[8] | IDH1, NPM1 mutated; DNMT3A, ZRSR2 gene variant | CD34, CD117, CD13, CD33, HLA-DR (partial), CD45. | AML-M4 |
| 189 | 86.94 | 64 | m | fav. | 82 | 75 | normal | NPM1, TET2 2 mutated; PHF6, BRAF gene variant | CD13, CD33, CD105 (partial), CD123, CD56, CD38, GlycophoinA (partial), cytoplasmatic TCL. | undifferentiated AML |
| 190 | 127.79 | 52 | m | int. | 72 | 75 | normal | NPM1, DNMT3A, SF3B1, TET2, FLT3 mutated | CD34, CD117, CD13/CD33, HLA-DR. | AML-M4 |

| | | | | | | | | | | |
|---|---|---|---|---|---|---|---|---|---|---|
| 191 | 103.76 | 46 | f | int. | 79.5 | 95 | normal | IDH2, NPM1, CBL mutated; FLT3 ITD | CD117, CD13/CD33, MPO. | AML-M1 |
| 192 | 34.70 | 60 | m | adv. | 22 | 90 | +8 | IDH2, RUNX1, SRSF2, DNMT3A, EED mutated | Myeloid population: CD34, CD117, CD13,HLA-DR+, partial MPO partial, CD105, CD123. Monocytic population: CD64, CD14, IREM2, strong aberrant expression of MPO and reduced expression of HLA-DR, CD35 and CD13. | AML-M1 |
| 193 | 603.41 | 37 | m | fav. | 72.5 | 90 | t(15;17)(q24;q21),t(16;18)(p?13;q?12) | PML-RARA | CD117, CD13, CD33, MPO, aberrant CD2, CD123. | AML-M3 |

**Table S2. Characteristics of AML patients.**

Age at diagnosis, sex, risk category, percentage of PB blasts and BM infiltration, cytogenetic aberration and molecular diagnosis, immunophenotype and FAB are listed for each patient for which RNA-Seq was performed.

Abbreviations: PB, peripheral blood, BM, bone marrow, adv., adverse; n.a., not available; MDS/MPS, myelodysplastic syndrome-myeloproliferative neoplasms.

| ID | Age at diagnosis | Sex | Risk | PB blasts (%) | BM infiltr. (%) | Cytogenetics | Molecular diagnosis | Immuno-phenotype | FAB |
|---|---|---|---|---|---|---|---|---|---|
| 1 | 54 | f | adv. | 85 | 90 | Deletion 7; EVI1 rearrangement | EVI1 positive: NRAS mutated | CD34, CD38, CD117, HLA-DR, CD13, CD33, CD4, CD7 (partial) and CD56 (partial). | secondary AML therapy related |
| 2 | 72 | m | n.a. | 20 | n.a. | n.a. | n.a. | two blast subpopulation: one with CD34, HLA-DR, CD33, CD71 and one with CD34, CD33, CD11b, CD35 and CD71. | secondary AML from MDS/MPS |
| 3 | 60 | f | adv. | 4.5 | 25 | Deletion (20p) | SF3B1 and ASXL1 mutated | | AML-M2 |
